# Early life starvation and Hedgehog-related signaling activate innate immunity downstream of *daf-18/PTEN* and *lin-35/Rb* causing developmental abnormalities in adult C. elegans

**DOI:** 10.1101/2025.02.28.640733

**Authors:** Ivan B. Falsztyn, Jingxian Chen, Rojin Chitrakar, James M. Jordan, L. Ryan Baugh

## Abstract

Early life experiences such as malnutrition can affect development and adult disease risk, but the molecular basis of such protracted effects is poorly understood. In the nematode *C. elegans,* extended starvation during the first larval stage causes the development of germline tumors and other abnormalities in the adult gonad, limiting reproductive success. Insulin/IGF signaling (IIS) acts through WNT signaling and lipid metabolism to promote starvation-induced gonad abnormalities, but IIS-independent modifiers have not been identified. We show that the tumor suppressors *daf-18/PTEN* and *lin-35/Rb* act independently of IIS to suppress starvation-induced abnormalities. We found that *lin-35/Rb* antagonizes activity of the Hedgehog (Hh) signaling homologs *ptr-23/PTCH-related*, *wrt-1/Hh-like*, *wrt-10/Hh-like*, and *tra-1/GLI*, which promote starvation-induced abnormalities. These Hh-related genes transcriptionally activate several genes associated with innate immunity in adults, which also promote starvation-induced gonad abnormalities. Surprisingly, we found that in addition to causing developmental abnormalities, early-life starvation induces an innate immune response later in life, leading to increased resistance to multiple bacterial pathogens. This work identifies a critical tumor-suppressor function of *daf-18/PTEN* independent of IIS, and it defines a regulatory network, including *lin-35/Rb*, Hh-related signaling, and the innate immunity pathway, that affects development of tumors and other developmental abnormalities resulting from early life starvation. By revealing that early-life starvation increases immunity later in life, this work suggests a fitness tradeoff between pathogen resistance and developmental robustness.

**AUTHOR SUMMARY:** Early life malnutrition can promote adult disease, including the risk of developing cancer. We previously found that the roundworm *Caenorhabditis elegans* develops tumors in well-fed adults following early life starvation, reflecting a breakdown of developmental fidelity. This powerful model system provides an excellent opportunity to identify molecular mechanisms that mediate the effects of early life starvation on development and adult physiology. Reducing insulin-like/IGF signaling during recovery from starvation suppresses tumor formation, but other regulatory pathways have not been identified. Here we show that a pair of important tumor suppressor genes, *daf-18/PTEN* and *lin-35/Rb*, function independently of insulin/IGF signaling to suppress starvation-induced abnormalities. *lin-35/Rb* represses the activity of genes involved in Hedgehog signaling, a cancer-causing pathway, which promote starvation-induced abnormalities. Hedgehog signaling activates an immune response, which, surprisingly, contributes to formation of tumors but renders adult worms resistant to pathogenic bacteria. Adult activation of this immune response by early life starvation suggests an evolutionary fitness tradeoff between pathogen resistance and developmental fidelity.

## INTRODUCTION

Fetal malnutrition is associated with increased risk of type II diabetes, cancer, and obesity in adulthood (Ravelli *et al*. 1976; Godfrey and Barker 2001; Walker and Ho 2012). Given confounding genetic and environmental factors in humans, tractable animal models are necessary to identify the molecular basis of such developmental origins of adult health and disease. The nematode *Caenorhabditis elegans* remains in a developmentally arrested state in the first larval stage (L1) after hatching without food (“L1 arrest” or “L1 diapause”) (Baugh 2013; Baugh and Hu 2020). Starved larvae are able to survive L1 arrest for a couple of weeks and resume development upon feeding. However, a significant fraction of individuals subjected to extended L1 arrest (*e.g.*, 8 days) develop proximal germline tumors, differentiated uterine masses, and other reproductive abnormalities on the first day of adulthood following recovery in replete conditions (Jordan *et al*. 2019, 2023; Shaul *et al*. 2023; Falsztyn *et al*. 2025). With its short generation time, ease of manipulation, and powerful genetic toolkit, *C. elegans* L1 arrest and recovery provides a powerful system to study the effects of early life starvation on adult health and disease.

Insulin/insulin-like growth factor (IGF) signaling (IIS) regulates developmental plasticity, aging, metabolism, and starvation resistance (Baugh 2013; Murphy and Hu 2013; Baugh and Hu 2020). Disruption of the sole known insulin/IGF receptor DAF-2/InsR increases survival during L1 arrest (Muñoz and Riddle 2003; Baugh and Sternberg 2006; Hibshman *et al*. 2017), and *daf-2/InsR* RNAi during larval development after L1 arrest suppresses starvation-induced gonad abnormalities, including germline tumors and uterine masses (Jordan *et al*. 2019, 2023; Shaul *et al*. 2023; Falsztyn *et al*. 2025). DAF-2/InsR signals through the phosphoinositide 3-kinase (PI3K) pathway by activating AGE-1/PI3K, which phosphorylates phosphatidylinositol-4, 5-bisphosphate (PIP2) to produce phosphatidylinositol-3, 4, 5-triphosphate (PIP3) (Dorman *et al*. 1995; Morris *et al*. 1996; Murphy and Hu 2013). PIP3 activates AKT and PDK kinases, which antagonize the forkhead box O transcription factor DAF-16/FoxO (Yen *et al*. 2011). Disruption of *daf-2/InsR* increases DAF-16/FoxO activity, and *daf-16* is required for starvation resistance and suppression of starvation-induced gonad abnormalities with *daf-2* RNAi (Baugh and Sternberg 2006; Jordan *et al*. 2019). WNT signaling and lipid metabolism operate downstream of IIS to promote starvation-induced abnormalities (Jordan *et al*. 2023; Shaul *et al*. 2023), but pathways affecting such abnormalities independently of IIS have not been identified.

The tumor suppressor phosphatase and tensin (PTEN) homolog DAF-18/PTEN inhibits PI3K/IIS by dephosphorylating PIP3 to produce PIP2, counteracting AGE-1/PI3K activity (Ogg and Ruvkun 1998). Loss of *daf-18/PTEN* constitutively activates IIS, inhibiting *daf-16/FoxO*, thereby reducing starvation resistance and enhancing development of starvation-induced gonad abnormalities (Fukuyama *et al*. 2012; Wolf *et al*. 2014; Shaul *et al*. 2023). However, disruption of *daf-2/InsR* does not completely suppress starvation-induced abnormalities, and loss of *daf-16/FoxO* does not increase penetrance of such abnormalities, though loss of *daf-18/PTEN* does. These observations suggest that DAF-18 functions independently of IIS to suppress development of germline tumors following early life starvation. In addition to being a lipid phosphatase, PTEN is a protein phosphatase (Myers *et al*. 1997), supporting the possibility that DAF-18/PTEN tumor-suppressor function may extend beyond dephosphorylation of PIP3 and inhibition of PI3K/IIS.

Here we show that *daf-18/PTEN* functions independently of IIS during larval development after L1 arrest to suppress starvation-induced gonad abnormalities. *daf-18* does so by acting through another important tumor suppressor, *lin-35/Rb*. We show that *daf-18* and *lin-35* inhibit activity of Hedgehog signaling homologs (Hh-related) to mediate suppression. Bulk RNA-seq and reporter gene analysis in adults subjected to extended L1 arrest suggests that Hh-related signaling promotes transcription of genes associated with innate immunity. These innate immunity genes function downstream of Hh-related signaling to promote starvation-induced abnormalities. Furthermore, we report that extended L1 arrest leads to induction of the innate immune pathway during late larval development without pathogen exposure, resulting in increased resistance to bacterial pathogens. This work is significant for implicating additional genes and pathways in development of tumors following early life starvation. This work is also important for revealing an apparent tension between developmental robustness and anticipation of pathogen exposure during recovery from starvation, adding an additional dimension to the complex interplay between development and the environment.

## RESULTS

### *daf-18/PTEN* suppresses starvation-induced gonad abnormalities independently of insulin/IGF signaling

*daf-18/PTEN* mutants are very sensitive to starvation and die rapidly during L1 arrest, but ethanol increases survival (Fukuyama *et al*. 2012; Chen *et al*. 2022). Wild-type larvae subjected to extended L1 arrest (*i.e*., 8 days) develop proximal germ cell tumors, differentiated uterine masses, and other gonad abnormalities (Jordan *et al*. 2019, 2023; Shaul *et al*. 2023; Falsztyn *et al*. 2025) as illustrated in Fig. S1A, B. With just 4 days of L1 starvation supplemented with ethanol, ∼50% of *daf-18(ok480)* null mutant individuals developed one or more gonad abnormalities compared to almost none of the wild type (Fig. 1A). After 8 days of starvation with ethanol, 100% of *daf-18* mutant worms developed abnormalities compared to ∼10% of wild type. These results show that *daf-18* is a potent suppressor of starvation-induced gonad abnormalities, including germ cell tumors and uterine masses. However, given that *daf-18* supports starvation survival, it was unclear from these results whether *daf-18* functions during L1 arrest and/or recovery to suppress abnormalities. We starved wild-type larvae for 8 days without ethanol and recovered them with *daf-18* RNAi bacteria. The frequency of starvation-induced abnormalities was significantly increased (Fig. 1B), showing that in addition to its role in starved L1 larvae, *daf-18* functions in fed, developing larvae to suppress starvation-induced abnormalities.

**Figure 1.**
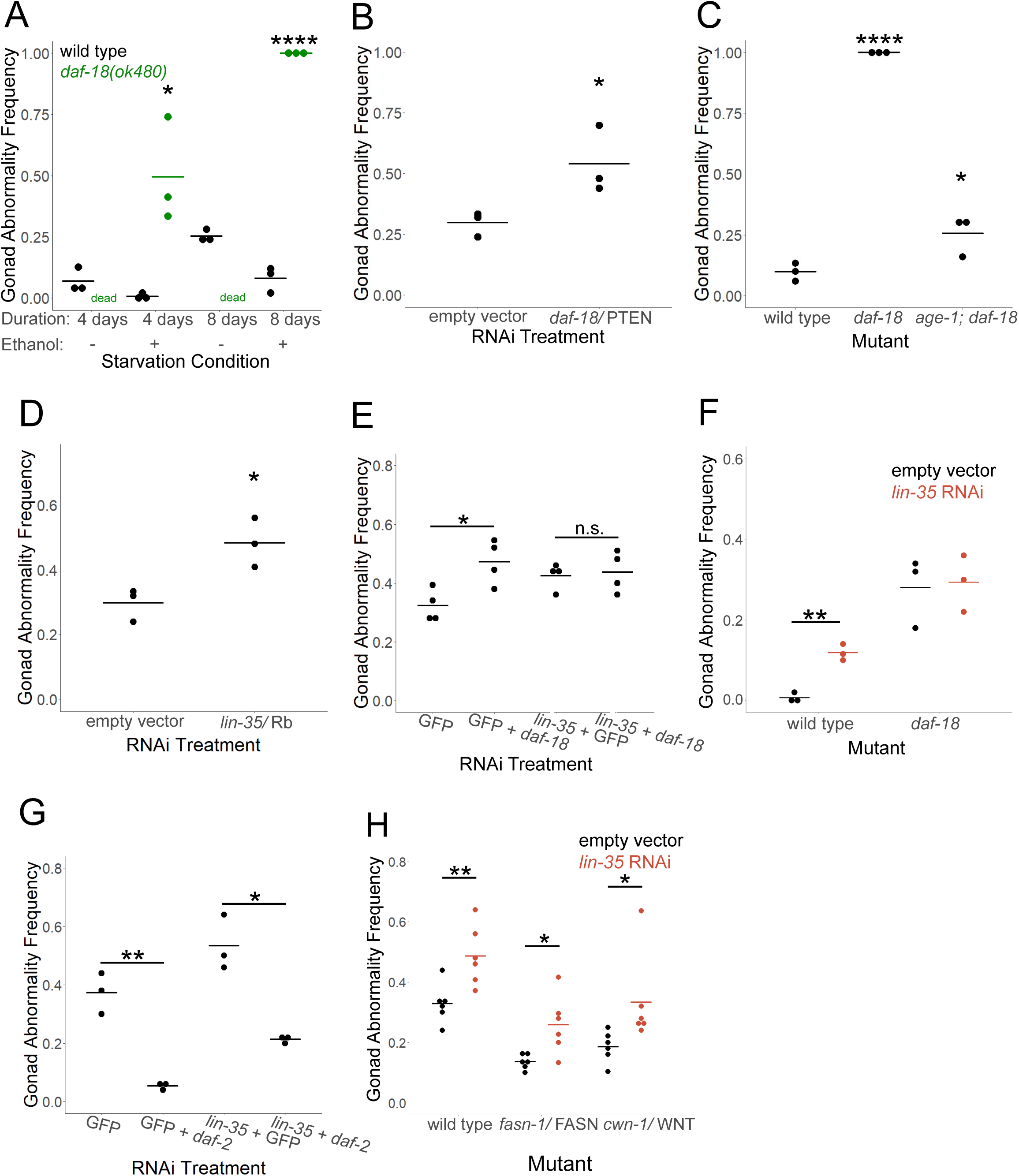
*daf-18/PTEN* and *lin-35/Rb* suppress starvation-induced gonad abnormalities independently of IIS. A-H) Gonad abnormalities (including proximal germ cell tumors, uterine masses, or other obvious developmental abnormalities in the gonad) were scored in various genotypes and conditions on the first day of egg laying. Starved worms were recovered with *E. coli* HT155 RNAi bacteria. A) Wild type and *daf-18(ok480)* mutants were starved for 4 or 8 days during L1 arrest in S-basal with or without 0.1% ethanol. *daf-18(ok480)* does not survive 4 or 8 days of starvation without ethanol (“dead”). B-E, G, H) Gonad abnormalities were scored after 8 days of L1 arrest. B) Wild-type worms were starved in virgin S-basal (no ethanol or cholesterol) and recovered on empty vector or *daf-18* RNAi bacteria. C) Wild-type, *daf-18(ok480)*, and *age-1(m333); daf-18(ok480)* worms were starved in S-basal with 0.1% ethanol. *daf-18* vs. *age-1; daf-18* p-value = 9.57e-05. D-E, G) Wild-type worms were starved in virgin S-basal and recovered on the indicated RNAi bacteria. E, G) GFP RNAi was used as a negative control for double-RNAi. F-G) Wild type and *daf-18(ok480)* (F) or wild type, *fasn-1(g14)*, and *cwn-1(ok546)* (G) were recovered with empty vector bacteria or *lin-35* RNAi bacteria. A-H) Each dot represents a biological replicate of ∼50 individuals (see Table S1 for summary statistics). Horizontal bars represent the mean across replicates. Asterisks indicate statistical significance compared to wild type within the same starvation condition (A), compared to empty vector (B, D), compared to wild type (C), and as indicated by horizontal bars (E-H). A-H) * P < 0.05, ** P < 0.01, **** P < 0.0001, “n.s.” = not significant; unpaired, two-tailed t-tests.

Disruption of *daf-2/InsR* suppresses starvation-induced gonad abnormalities (Fig. S1C) (Jordan *et al*. 2019, 2023; Shaul *et al*. 2023; Falsztyn *et al*. 2025), and we wondered if the potent effect of *daf-18/PTEN* on such abnormalities is due to its inhibition of IIS. We confirmed (Jordan *et al*. 2019) that the transcription factors *daf-16/FoxO* and *skn-1/Nrf*, effectors of IIS, are required for *daf-2/InsR* RNAi to suppress starvation-induced abnormalities (Fig. S1C). However, disruption of *daf-16* and *skn-1* alone or together did not significantly increase the frequency of abnormalities, though *daf-18* RNAi did (Fig. 1B). Furthermore, a gain-of-function allele of *skn-1* and over-expression of *daf-16* incompletely suppressed abnormalities (Fig. S1D). DAF-16 and PQM-1 engage in mutual antagonism (Tepper *et al*. 2013), and PQM-1 could function as an additional transcriptional effector of IIS. However, *pqm-1* RNAi did not significantly affect development of gonad abnormalities in previously starved worms (Fig. S1E). Together these results suggest that changes in activity of the known effectors of IIS cannot account for the full effect of disruption of *daf-18/PTEN* on starvation-induced abnormalities.

PTEN dephosphorylates PIP3 to produce PIP2, inhibiting PI3K/IIS in mammals and *C. elegans*, but it can also dephosphorylate proteins (Myers *et al*. 1997; Ogg and Ruvkun 1998). PIP3 is not detectable in an *age-1/PI3K* null mutant (Bharill *et al*. 2013), so if the only function of DAF-18 in suppressing starvation-induced abnormalities is to inhibit PI3K signaling via its lipid-phosphatase activity, then loss of *age-1/PI3K* should rescue the enhanced abnormality phenotype of *daf-18(ok480)* to wild-type penetrance or less. The *age-1(m333); daf-18(ok480)* double null mutant displayed a significantly lower gonad abnormality frequency than *daf-18(ok480)*, reflecting inhibition of PI3K/IIS via its lipid-phosphatase activity. However, suppression was incomplete, and gonad abnormalities occurred at a significantly higher frequency in *age-1(m333); daf-18(ok480)* than wild type following starvation (Fig. 1C). This result suggests that in addition to inhibiting PI3K/IIS, DAF-18/PTEN suppresses starvation-induced gonad abnormalities independently of PI3K/IIS, potentially via its protein-phosphatase activity.

### *daf-18/PTEN* functions through *lin-35/Rb* to suppress starvation-induced gonad abnormalities

We recently discovered that *daf-18/PTEN* functions through *lin-35/Rb* to promote starvation survival during L1 arrest (Chen *et al*. 2025). LIN-35 is the sole worm homolog of the retinoblastoma (RB) pocket protein family (Lu and Robert Horvitz 1998). RB proteins are tumor suppressors (Berry *et al*. 2019), and we hypothesized that LIN-35/Rb suppresses formation of the proximal germ cell tumors and differentiated uterine masses which largely comprise the starvation-induced gonad abnormality phenotype. *lin-35* is required for vulva development (Lu and Robert Horvitz 1998). Without L1 starvation, *lin-35(n745)* null mutant larvae develop gonad abnormalities and other defects, and they often do not survive to adulthood, so we did not analyze them. Instead, we recovered wild-type worms subjected to 8 days of L1 arrest with RNAi bacteria targeting *lin-35*, which significantly increased the frequency of starvation-induced abnormalities (Fig. 1D). Notably, *daf-18* and *lin-35* RNAi did not cause gonad abnormalities without L1 starvation (Fig. S1F), demonstrating that abnormalities depend on starvation and do not result from general disruption of development. These results show that *lin-35* functions in fed, developing larvae to suppress starvation-induced abnormalities, like *daf-18* (Fig. 1B), suggesting *lin-35* could mediate the hypothetical PI3K/IIS-independent effect of *daf-18* on gonad abnormalities. Consistent with this hypothesis, disruption of *daf-18* and *lin-35* together did not increase the frequency of abnormalities compared to *lin-35* alone (Fig. 1E). This result was corroborated with *lin-35* RNAi and *daf-18(ok480)* (Fig. 1F). *daf-2/InsR* RNAi suppressed starvation-induced abnormalities (Fig. 1G), as expected (Jordan *et al*. 2019; Shaul *et al*. 2023; Falsztyn *et al*. 2025). However, disrupting *daf-2/InsR* and *lin-35/Rb* together suppressed abnormality frequency compared to *lin-35* alone (Fig. 1G), suggesting independent effects of IIS and *lin-35/Rb* on development of starvation-induced abnormalities. The sole fatty acid synthetase *fasn-1/FAS* and WNT signaling function downstream of IIS to promote starvation-induced abnormalities (Jordan *et al*. 2023; Shaul *et al*. 2023). *lin-35* RNAi significantly increased the frequency of abnormalities in *fasn-1/FAS* and *cwn-1/WNT* mutant backgrounds (Fig. 1H), consistent with independent function of *lin-35*. Together these results suggest that *lin-35/Rb* functions downstream of *daf-18/PTEN* and independently of PI3K/IIS to suppress starvation-induced gonad abnormalities.

### Hh-related signaling promotes starvation-induced gonad abnormalities downstream of daf-18/PTEN *and* lin-35/Rb

*daf-2/InsR*, *daf-16/FoxO*, *daf-18/PTEN*, *lin-35/Rb*, *fasn-1/FAS*, and WNT signaling all modulate formation of starvation-induced gonad abnormalities, and their homologs affect tumor development in mammals, suggesting that homologs of other genes that promote or suppress tumor formation could affect starvation-induced abnormalities. Like WNT signaling, Hedgehog (Hh) signaling promotes stem cell proliferation and tumor formation (Nüsslein-Volhard and Wieschaus 1980; Di Magliano and Hebrok 2003; Jiang and Hui 2008), and we hypothesized that Hh signaling promotes starvation-induced gonad abnormalities. Most genes involved Hh signaling have many homologs in *C. elegans*, some homologs are missing (*e.g.*, Smoothened), and in some cases there are missing or additional protein domains (Aspöck *et al*. 1999). Consequently, it is relatively mysterious how Hh-related signaling works in *C. elegans*, though a variety of phenotypes related to molting, reproductive aging, germ cell proliferation, and innate immunity have been reported (Kuwabara *et al*. 2000; Hao *et al*. 2006; Templeman *et al*. 2020; Zárate-Potes *et al*. 2022; Wang *et al*. 2023). Nonetheless, we found that mutation of the PATCHED/PTCH receptor-related gene *ptr-23* and the Hedgehog-related genes *wrt-1* and *wrt-10* suppressed starvation-induced gonad abnormalities (Fig. 2A). *ptr-23*, *wrt-1*, and *wrt-10* mutants did not survive longer than wild type during L1 arrest (Fig. S2A), suggesting the decrease in abnormality frequency is not due to a general increase in starvation resistance in the mutants. RNAi of these genes during recovery from L1 arrest also suppressed starvation-induced abnormalities (Fig. 2B), suggesting that Hh-related signaling during larval development promotes formation of starvation-induced gonad abnormalities.

**Figure 2.**
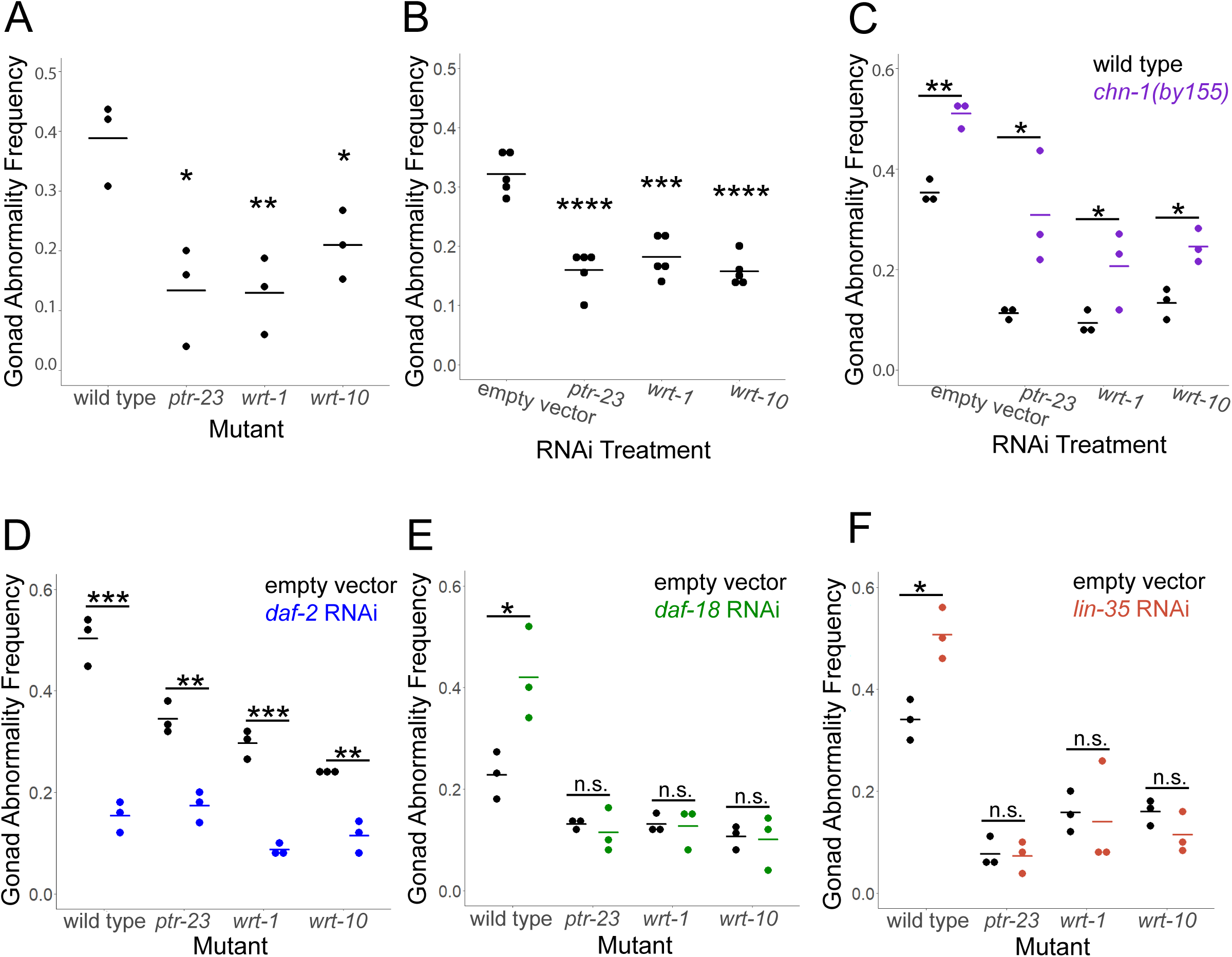
Hh-related genes promote starvation-induced gonad abnormalities downstream of *daf-18/PTEN* and *lin-35/Rb.* A-F) Larvae were starved in L1 arrest for 8 days in virgin S-basal (S-basal without ethanol or cholesterol), and gonad abnormalities were scored. A) Wild type and *ptr-23(ok3663)*, *wrt-1(tm1417)*, and *wrt-10(aus36)*) were recovered on empty vector RNAi bacteria. B) Wild-type larvae were recovered on empty vector RNAi bacteria and RNAi targeting *ptr-23*, *wrt-1*, and *wrt-10*. C) Wild type and *chn-1(by155)* were recovered on empty vector RNAi bacteria and RNAi bacteria targeting *ptr-23*, *wrt-1*, and *wrt-10*. D-F) Wild type and *ptr-23*, *wrt-1*, and *wrt-10* mutants were recovered on empty vector RNAi bacteria and *daf-2* RNAi (D), *daf-18* RNAi (E), and *lin-35* RNAi (F). A-F) Each dot represents a biological replicate with ∼50 individuals (see Table S1 for summary statistics). Horizontal bars represent the mean across replicates. Asterisks indicate statistical significance relative to wild type (A), relative to empty vector RNAi (B), and as indicated by bars (C-F). * P < 0.05, ** P < 0.01, *** P < 0.001, **** P < 0.0001, “n.s.” = not significant; unpaired, two-tailed t-test.

Having established a role for Hh-related signaling in development of starvation-induced gonad abnormalities, we asked whether Hh-related signaling functions independently of IIS. The ubiquitin ligase CHN-1/CHIP ubiquitinates DAF-2/InsR promoting its degradation, and loss of *chn-1/CHIP* increases DAF-2 abundance and IIS (Tawo *et al*. 2017). The *chn-1(by155)* null mutant increased the frequency of starvation-induced abnormalities (Fig. 2C), as expected (Falsztyn *et al*. 2025). The mutant also increased abnormality frequency along with RNAi targeting *ptr-23*, *wrt-1*, and *wrt-10*, suggesting that Hh-related signaling functions independently of IIS. *chn-1* mutant larvae did not develop gonad abnormalities without extended L1 arrest (Fig. S2B), demonstrating that abnormalities depend on starvation and do not result from general disruption of development. CHN-1 may have targets in addition to DAF-2. However, *daf-2* RNAi suppressed abnormality frequency in *ptr-23*, *wrt-1*, and *wrt-10* mutants (Fig. 2D), supporting the conclusion that Hh-related signaling functions independently of IIS to promote starvation-induced gonad abnormalities.

Given that Hh-related signaling functions independently of IIS in this context, we hypothesized that it functions downstream of *daf-18/PTEN* and *lin-35/Rb*. *daf-18* RNAi and *lin-35* RNAi did not increase penetrance of starvation-induced abnormalities in *ptr-23*, *wrt-1*, or *wrt-10* mutant backgrounds (Fig. 2E, F), suggesting that these Hh-related signaling genes are epistatic to *daf-18* and *lin-35*. These results support the conclusion that Hh-related signaling promotes starvation-induced gonad abnormalities downstream of *daf-18/PTEN* and *lin-35/Rb*.

### Hh-related genes function in the hypodermis and intestine to promote starvation-induced gonad abnormalities

We made transcriptional reporter genes for *ptr-23*, *wrt-1*, and *wrt-10* to determine where they are expressed during larval development and to investigate regulation of their expression. Each reporter gene was expressed throughout the hypodermis (Fig. 3A), consistent with single-cell mRNA sequencing (scRNA-seq) of L2 larvae (Cao *et al*. 2017) and reporter gene analysis of *ptr-23* (Rohlfing *et al*. 2011). Each reporter gene was also expressed throughout the intestine, though only *ptr-23* had substantial expression there according to scRNA-seq (Cao *et al*. 2017). Dim expression was also observed in the pharynx for *ptr-23p::YFP* and *wrt-10p::YFP*, and in the vulva for *ptr-23p::YFP*, consistent with scRNA-seq (Cao *et al*. 2017). Additional unidentified cells in the head and tail were also observed.

**Figure 3.**
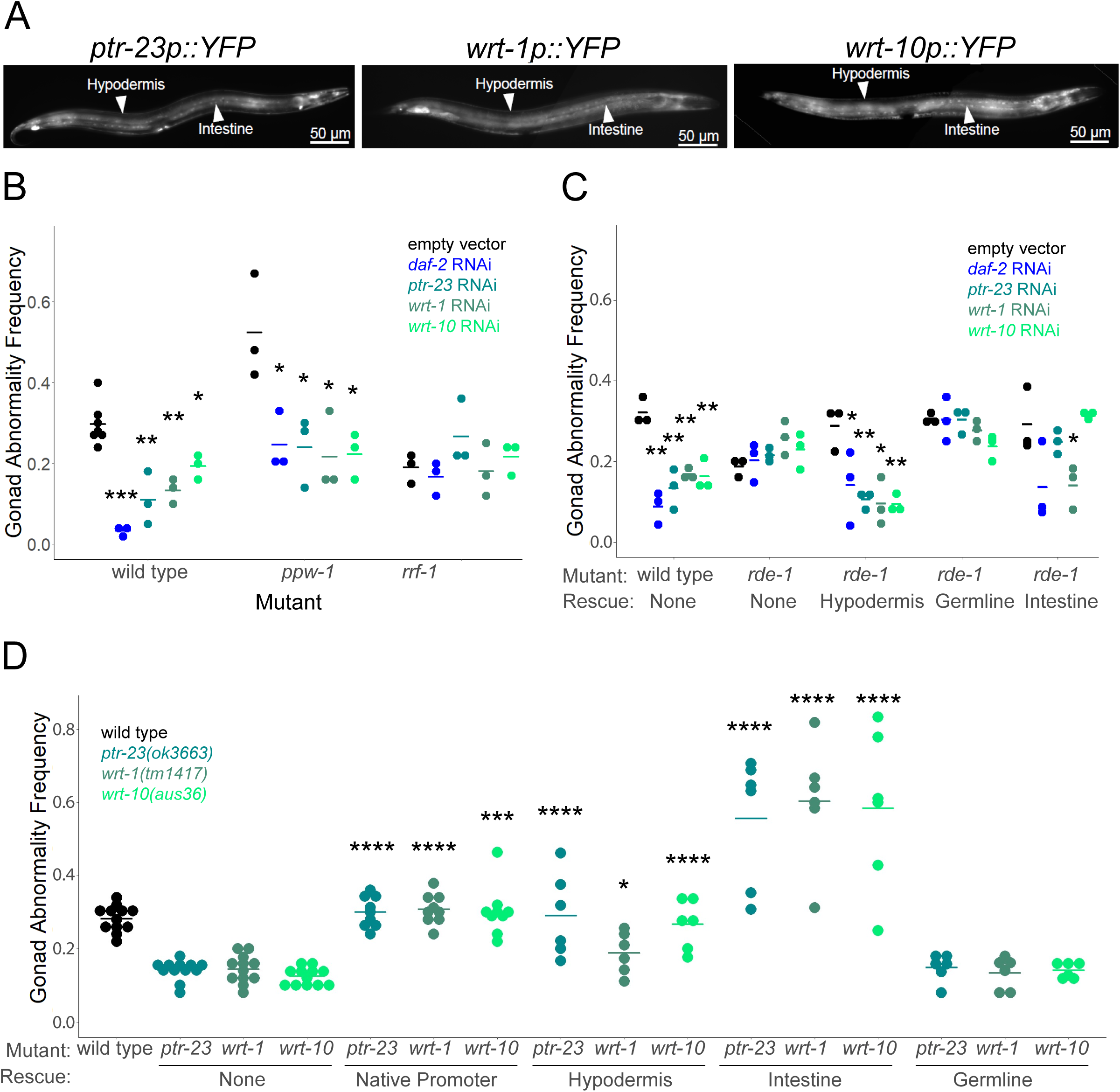
Hh-related genes function in the hypodermis and intestine to promote starvation-induced gonad abnormalities. A) Representative images of *ptr-23p::YFP, wrt-1p::YFP,* and *wrt-10p::YFP* at 400X total magnification show expression primarily in the hypodermis and intestine as indicated with arrowheads and labels. Images were taken in L4 larvae (48 hours after hatching) in well-fed individuals. B) Wild type, *ppw-1(pk2505),* and *rrf-1(pk1417)* L1 larvae were arrested for 8 days as L1 larvae then recovered on empty vector, *daf-2*, *ptr-23, wrt-1,* or *wrt-10* RNAi bacteria and scored for gonad abnormalities. C) Wild type and *rde-1(ne219)* mutants were starved for 8 days then recovered on empty vector, *ptr-23, wrt-1,* and *wrt-10* RNAi and scored for gonad abnormalities. *rde-1(ne219)* null mutants that had been rescued with a multicopy *rde-1* transgene with tissue-specific expression in the hypodermis (*lin-26p*), germline (*mex-5p*), or intestine (*nhx-2*) were also included. D) Wild type and *ptr-23(ok3663), wrt-1(tm1417),* and *wrt-10(au36)* were starved for 8 days as L1 larvae then recovered on empty vector RNAi bacteria and scored for gonad abnormalities. *ptr-23(ok3663), wrt-1(tm1417),* and *wrt-10(au36)* mutants that had been rescued with a multicopy transgene with their native promoter or a tissue-specific promoter for the hypodermis (*col-12p*), intestine (*ges-1p*), or germline (*mex-5p*), were also scored. Gonad abnormalities are significantly greater than wild type with intestinal rescue of each Hh-related gene (P < 0.0001). B-D) Each dot represents a biological replicate with ∼50 individuals (see Table S1 for summary statistics). Horizontal bars represent the mean across replicates. Asterisks indicate significance between the indicated RNAi treatment and empty vector RNAi bacteria within the same genotype (B-C), and between the indicated mutants with transgenic rescue and the corresponding mutant without rescue (“none”) (D). * P < 0.05, ** P < 0.01, *** P < 0.001, **** P < 0.0001; Unpaired, two-tailed t-test.

Whole-worm, image-based quantification of reporter gene expression revealed no effect of 8 days L1 arrest on subsequent expression of the reporters (Fig. S3A). These results suggest that in general L1 starvation does not promote gonad abnormalities by increasing transcription of Hh-related genes later in life. Image-based analysis also did not reveal an effect of *daf-18/PTEN* or *lin-35/Rb* RNAi on reporter expression levels, with the exception of the *ptr-23p::YFP* reporter after 1 day of L1 arrest (Fig. S3A). Nonetheless, it is possible that multicopy reporters lack the sensitivity to capture significant changes in gene expression.

We used multiple complementary approaches to determine the anatomical site(s) of action for *ptr-23*, *wrt-1*, and *wrt-10* in promoting starvation-induced gonad abnormalities. *ppw-1* and *rrf-1* mutants are generally deficient for germline and somatic RNAi, respectively (Sijen *et al*. 2001; Tijsterman *et al*. 2002). *daf-2/InsR* RNAi suppressed abnormalities in wild-type and *ppw-1* mutants but not *rrf-1* (Fig. 3B), as expected (Jordan *et al*. 2019). RNAi is partially effective in some somatic tissues in *rrf-1* mutants (Kumsta and Hansen 2012), but this result nonetheless suggests that *daf-2* functions in the soma, as opposed to the germline, to suppress abnormalities, as shown elsewhere (Jordan *et al*. 2019). Likewise, RNAi of *ptr-23*, *wrt-1*, and *wrt-10* suppressed abnormalities in wild-type and *ppw-1* mutant backgrounds but not the *rrf-1* background (Fig. 3B), suggesting that Hh-related signaling functions in the soma to promote gonad abnormalities.

*rde-1* is required for RNAi throughout the animal (Tabara *et al*. 1999), and we used strains with tissue-specific transgenic rescue of an *rde-1* mutation to further evaluate potential site(s) of action of *ptr-23*, *wrt-1*, and *wrt-10.* Mutation of *rde-1* abrogated the effects of RNAi for *daf-2* and each of the Hh-related genes (Fig. 3C), as expected. Rescue of *rde-1* in the hypodermis (*lin-26p*, (Qadota *et al*. 2007) restored suppression of starvation induced-abnormalities following *ptr-23*, *wrt-1*, and *wrt-10* RNAi (Fig. 3C), consistent with prominent expression of all three genes in the hypodermis (Cao *et al*. 2017). Intestinal rescue (*nhx-2p*, (Espelt *et al*. 2005) produced mixed results, with *wrt-1* RNAi suppressing abnormalities but not *ptr-23* or *wrt-10.* Germline rescue (*mex-5p*, (Marré *et al*. 2016) of *rde-1* did not result in suppression of abnormalities with RNAi of any gene tested (Fig. 3C), consistent with analysis of *ppw-1* and *rrf-1* mutants (Fig. 3B). These results suggest that all three Hh-related genes function in the hypodermis and that *wrt-10* also functions in the intestine to suppress abnormalities.

Transgenic rescue of *ptr-23*, *wrt-1*, and *wrt-10* with their own promoters restored wild-type penetrance of gonad abnormalities (Fig. 3D), as expected. Hypodermal rescue (*col-12p*) of each gene also increased the penetrance of starvation-induced abnormalities, but germline rescue (*mex-5p*) did not, consistent with tissue-specific RNAi (Fig. 3B, C). Surprisingly, intestinal rescue (*ges-1p*) of all three genes increased penetrance above wild-type levels (Fig. 3D), likely due in part to over-expression with a heterologous promoter and multi-copy transgene. Intestinal rescue did not cause gonad abnormalities without extended L1 arrest (Fig. S3C), demonstrating that abnormalities depend on starvation and do not result from general disruption of development. Together with our tissue-specific RNAi results, these results suggest that *wrt-1* is necessary and sufficient in both hypodermis and intestine to promote starvation-induced gonad abnormalities, and that *ptr-23* and *wrt-10* are necessary and sufficient in the hypodermis but merely sufficient in the intestine to do so. Moreover, these results suggest cell-nonautonomous effects of Hh-related signaling on germ cell proliferation and other aspects of gonad development affected by extended L1 arrest.

### Hh-related genes promote expression of genes related to innate immunity

We used RNA sequencing (RNA-seq) of whole adult worms to identify genes whose expression is affected by Hh-related signaling following extended L1 arrest. We starved wild-type L1 larvae for 8 days, recovered them with empty vector RNAi bacteria or RNAi targeting *ptr-23*, *wrt-1*, *wrt-10*, or *tra-1/GLI*, collected them on the first day of egg laying, and prepared RNA for sequencing. *tra-1* encodes the sole worm homolog of the GLI transcription factor family (Mathies *et al*. 2004). GLI factors function as transcriptional effectors of canonical Hh signaling (Sigafoos *et al*. 2021). Although *tra-1/GLI* has not been shown to function in Hh-related signaling, we included it in our RNA-seq experiment since it was unclear how the other Hh-related genes could affect transcription. Notably, *tra-1* RNAi suppressed starvation-induced abnormalities (Fig. S4), suggesting it could mediate effects of *ptr-23*, *wrt-1*, and *wrt-10.* Cluster analysis of 365 genes differentially expressed across the entire set of conditions (generalized linear model) suggested that all four Hh-related genes had similar effects on gene expression (Fig. 4A). Relatively few genes were significantly differentially expressed in any one perturbation relative to the control (exact test), but a majority were affected by three or four of the perturbations (Fig. 4B). Together these results suggest that the Hh-related genes converge on transcriptional regulation of a common set of genes. We treated all replicates for all four RNAi treatments as a single perturbation to identify additional genes affected by Hh-related signaling. Strikingly, 22 of the 41 genes downregulated by Hh-related RNAi are associated with innate immunity. Fourteen genes are categorized as ‘Stress response: pathogen’ (p-value = 1.65E-18) and eight are categorized as ‘Stress response: C-type Lectin’ (p-value = 9.96E-08), which are associated with innate immunity (Holdorf *et al*. 2020; Liu *et al*. 2024), but none of the 19 upregulated genes are associated with innate immunity (Fig. 4C). These gene expression results suggest that Hh-related signaling promotes expression of genes related to innate immunity.

**Figure 4.**
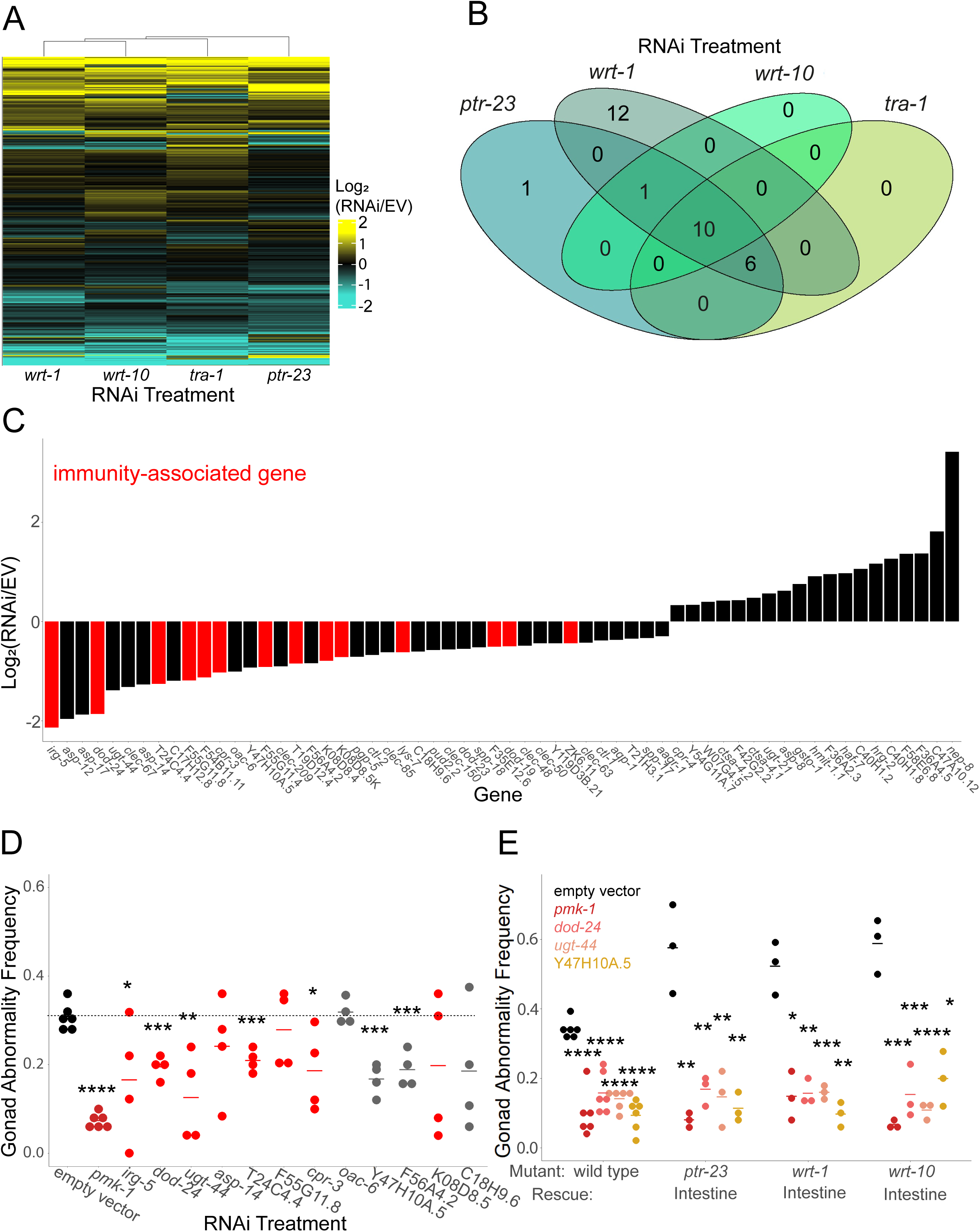
Hh-related signaling promotes innate immunity gene expression to promote starvation-induced gonad abnormalities. A) RNA-seq was performed on the first day of egg laying in worms that had been subjected to 8 days of starvation during L1 arrest and then recovered with empty vector, *ptr-23, tra-1, wrt-1,* or *wrt-10* RNAi bacteria. Log2 fold-changes of transcript abundance between RNAi of each gene and empty vector were clustered and are presented as a heat map. Each horizontal bar represents expression of a gene. A generalized linear model was used to identify differentially expressed genes across all four treatments (P < 0.05), and these genes were used for cluster analysis (n = 365). B) The number of genes that were differentially expressed in each pairwise comparison between RNAi and empty vector is plotted in a Venn diagram. An exact test (EdgeR) was used to identify differentially expressed genes (FDR < 0.1). C) All replicates for all RNAi treatments were combined and treated as a single perturbation to compare to empty vector to identify a larger set of differentially expressed genes with the exact test. The resulting list of differentially expressed genes (FDR < 0.1) was sorted by fold change, and log_2_ fold-change is plotted. Genes identified by WormCat as components of the innate immune system are highlighted red. D) Wild type L1 larvae were starved for 8 days then recovered with empty vector RNAi bacteria, *pmk-1* RNAi (positive control), or RNAi against twelve of the down-regulated genes plotted in C, then scored for gonad abnormalities on the first day of egg laying. Genes identified by WormCat as components of the innate immune system are in red. The four genes in gray were upregulated by pathogen exposure and downregulated by loss of *pmk-1/p38 MAPK* (Fletcher *et al*. 2019). E) Wild type and *ptr-23(ok3663), wrt-1(tm1417),* and *wrt-10(au36)* with intestinal rescue (Fig. 3D) were starved for 8 days as L1 larvae and recovered with empty vector RNAi bacteria, *pmk-1* RNAi, or RNAi against three of the genes analyzed in D. D-E) Each dot represents a biological replicate with ∼50 individuals (see Table S1 for summary statistics). Horizontal bars represent the mean across replicates. Asterisks indicate significance between the indicated RNAi treatment and empty vector RNAi (D) or empty vector RNAi within the same genotype (E). * P < 0.05, ** P < 0.01, *** P < 0.001, **** P < 0.0001; unpaired, two-tailed t-test.

### Innate immunity genes promote starvation-induced gonad abnormalities downstream of Hh-related signaling

We hypothesized that Hh-related signaling promotes expression of genes, specifically innate immunity genes, that promote starvation-induced gonad abnormalities. PMK-1/p38 MAPK is an essential component of the pathway that activates the innate immunity response to pathogens (Shivers *et al*. 2008). We targeted *pmk-1* with RNAi along with eight genes downregulated by Hh-related RNAi and associated with innate immunity. We also tested four downregulated genes that are not in WormCat innate immunity-related categories but were transcriptionally upregulated upon pathogen exposure and downregulated in *pmk-*1 null mutants in another study (Fletcher *et al*. 2019). RNAi of *pmk-1* and seven of the other twelve genes tested significantly decreased the frequency of starvation-induced abnormalities (Fig. 4D). These results suggest that genes associated with the innate immunity pathway, including its critical regulator *pmk-1/p38 MAPK*, promotes starvation-induced gonad abnormalities. We took advantage of the elevated frequency of starvation-induced abnormalities resulting from intestinal overexpression of *ptr-23*, *wrt-1*, and *wrt-10* (Fig. 3D) to determine if these immunity-related genes function downstream of Hh-related signaling. RNAi of *pmk-1* and the three other genes tested suppressed gonad abnormalities despite intestinal overexpression of *ptr-23*, *wrt-1*, and *wrt-10* (Fig. 3E). These results suggest the innate immunity pathway, or genes associated with it, promotes starvation-induced gonad abnormalities downstream of Hh-related signaling.

### Starvation during L1 arrest induces the innate immunity pathway and pathogen resistance later in life

We were surprised to find that genes associated with innate immunity promote starvation-induced abnormalities, and we wondered if they actually function in the innate immunity pathway to do so. We reasoned that if they do, then starvation during L1 arrest should result in increased resistance to bacterial pathogens. Remarkably, L4 larvae starved for 1 or 8 days during L1 arrest displayed increased resistance to *P. aeruginosa* strain PA14 in a “fast-killing” assay (Tan *et al*. 1999) compared to control worms that had not been starved during L1 arrest (“unstarved”), with 8 days of starvation having a larger effect than 1 day (Fig. 5A). Moreover, L4 larvae starved for 8 days during L1 arrest displayed increased resistance to PA14 in a “slow-killing” assay (Tan *et al*. 1999) and to *S. enterica* (Fig. S5A, B). 1 or 8 days of L1 arrest was also sufficient to increase expression of a *sysm-1p::GFP* reporter gene in L4 larvae without pathogen exposure, with 8 days of starvation having a larger effect than 1 day (Fig. 5B and Fig. S5C). This result shows that *sysm-1*, a marker of the innate immune response (Shivers *et al*. 2009), is activated in L4 larvae by L1 starvation. To determine when the *sysm-1p::GFP* reporter is induced (*e.g.*, during starvation or recovery), we measured *sysm-1p::GFP* intensity in arrested L1 larvae and at different timepoints during recovery (Fig. 5C). There was no difference in intensity in L1 larvae that had been starved for 1 or 8 d, suggesting that reporter gene induction does not occur as a direct response to starvation. There was a modest but significant increase in expression after 6 hr recovery and an apparent increase over developmental time for larvae recovering from L1 arrest as well as unstarved controls. These observations suggest that exposure to food (non-pathogenic *E. coli*), provokes expression of *sysm-1p::GFP*. Notably, expression of *sysm-1p::GFP* was significantly lower in previously starved larvae than unstarved controls at 6 hr, and it was equivalent between them at 24 hr. However, *sysm-1p::GFP* expression was significantly higher at 40 hr for larvae that had been starved for 8 d compared to unstarved controls, and it was significantly higher for larvae that had been starved for 1 and 8 d at 48 hr, as expected (Fig. 5B). These results suggest that previously starved larvae require a substantial amount of recovery time for innate immunity pathway induction to exceed that of unstarved controls. Together these results show that early life starvation during L1 arrest activates innate immunity later in life, with a longer duration of starvation resulting in stronger activation.

**Figure 5.**
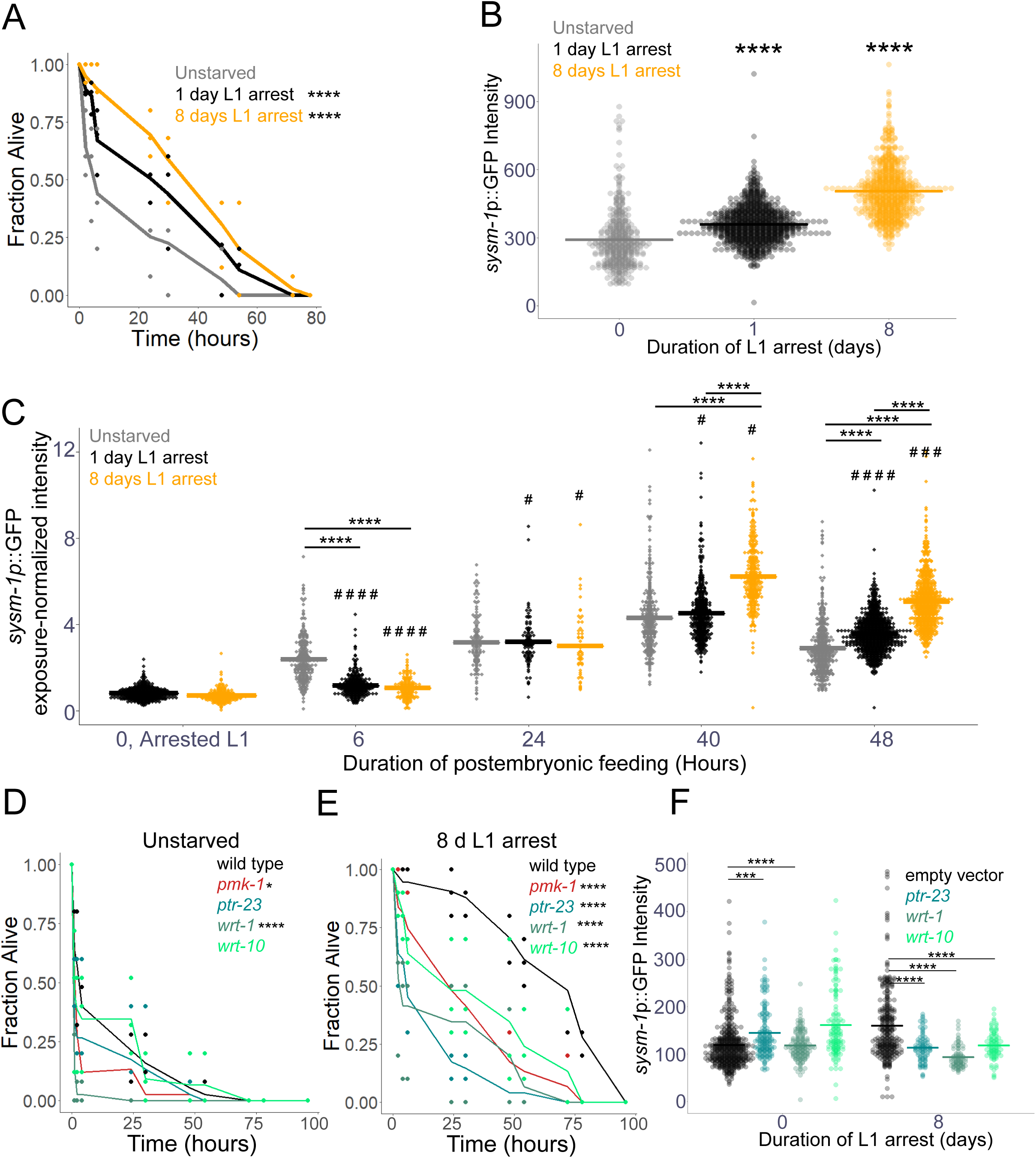
Starvation during L1 arrest promotes innate immunity later in life via Hh-related signaling. A) Wild-type L1 larvae were starved for the indicated duration, recovered on plates with *E. coli* HT115 bacteria, L4 larvae were transferred to *P. aeruginosa* PA14 “fast-killing” plates, and survival was assayed. B-C) L1 larvae with a *sysm-1p*::*GFP* reporter gene were starved for the indicated duration, recovered on plates with empty vector RNAi bacteria for 48 hr (B), or the indicated duration (C) and whole-worm reporter gene expression was quantified by image analysis. C) Due to varying exposure times, exposure-normalized whole worm GFP intensity (fluorescence intensity divided by exposure time) is reported. Pound signs indicate significant differences between 1 or 8 d L1 arrest and unstarved controls for a given duration of postembryonic feeding; # P < 0.05, ### P < 0.001, #### P < 0.0001. D, E) Unstarved worms (D) or worms that had been starved for 8 days as L1 larvae (E) were exposed to PA14 as in A, except that in addition to wild type, *pmk-1(km25)*, *ptr-23(ok3663)*, *wrt-1(tm1417)*, and *wrt-10(aus36)* were included. F) L1 larvae with a *sysm-1p*::*GFP* reporter gene were starved for 0 (unstarved) or 8 days, recovered, and imaged as in B, except that they were recovered with empty vector RNAi bacteria or RNAi targeting *ptr-23*, *wrt-1*, or *wrt-10.* B, F) To ensure matching stages, unstarved control worms (0 hr) were allowed 8 additional hours to account for embryogenesis, and worms starved for 8 days were also allowed 8 additional hours to account for developmental delay following extended starvation. A, D, E) Three biological replicates were performed with ∼50 individuals each (see Table S1 for summary statistics). Mean survival at each timepoint is plotted as a line with survival of each replicate at each timepoint included as dots. Asterisks indicate significance between the indicated starvation durations (1 or 8 d L1 arrest) and unstarved controls (A) or between the indicated genotype and wild type (D, E). The log-rank test was used for pairwise comparisons. B, C, F) Individual points represent the average, background-corrected pixel intensity for a single worm. Horizontal bars represent the mean intensity across three biological replicates. A linear mixed-effect model was fit to the data with background-corrected average intensity as the response variable, RNAi treatment as the fixed effect, and replicate as the random effect. Asterisks and bars indicate significance. A-F) * P < 0.05, *** P < 0.001, **** P < 0.0001.

### Hh-related signaling is required for L1 arrest to induce the innate immunity pathway

Since Hh-related signaling promotes starvation-induced gonad abnormalities (Fig. 2), innate immunity genes promote abnormalities downstream of Hh-related signaling (Fig. 4), and L1 starvation induces innate immunity (Fig. 5A, B), we hypothesized that Hh-related signaling promotes innate immunity in response to L1 starvation. Mutations disrupting function of *pmk-1/p38 MAPK* and *wrt-1*, but not *ptr-23* or *wrt-10*, significantly decreased resistance to PA14 in L4 larvae that had not been subjected to L1 starvation (Fig. 5C). In contrast, mutations affecting all four genes dramatically decreased resistance to PA14 in larvae that had been starved for 8 days during L1 arrest (Fig. 5D). Critically, the increase in resistance caused by extended L1 arrest was largely dependent on the Hh-related genes, with those mutants displaying pathogen sensitivity similar to *pmk-1*. Furthermore, RNAi of *ptr-23*, *wrt-1*, and *wrt-10* prevented activation of *sysm-1p::GFP* by 8 days of L1 arrest (Fig. 5E). Together these results suggest that induction of innate immunity in L4 larvae and adults by L1 starvation depends on Hh-related signaling.

## DISCUSSION

*C. elegans* L1 arrest and recovery provides a powerful system to investigate the molecular basis of adult consequences of early life starvation. We show that *daf-18/PTEN* suppresses L1 starvation-induced germline tumors and other gonad abnormalities in adults via its well-characterized lipid-phosphatase activity inhibiting PI3K/IIS (Fig. 6). In addition, we show that *daf-18* suppresses such starvation-induced abnormalities independently of PI3K/IIS, likely via its putative protein-phosphatase activity. We show that *lin-35/Rb* also suppresses starvation-induced abnormalities, and that it functions downstream of *daf-18* and independently of IIS. We identified three Hedgehog signaling homologs (*ptr-23/PTCH-related*, *wrt-1/Hh-related*, and *wrt-10/Hh-related*) that promote starvation-induced abnormalities and are antagonized by *lin-35*. Along with *tra-1/GLI*, a putative transcriptional effector of Hh-related signaling, these Hh-related genes affect expression of a common set of genes involved in the innate immunity pathway. We demonstrate that innate immunity genes promote development of starvation-induced abnormalities, and that L1 starvation induces immunity, causing increased pathogen resistance in L4 larvae. Induction of immunity late in development by early life starvation depends on Hh-related signaling, suggesting a potentially adaptive example of phenotypic plasticity regulated by a novel signaling pathway involving DAF-18/PTEN, LIN-35/Rb, Hh signaling homologs, and the innate immunity response.

**Figure 6.**
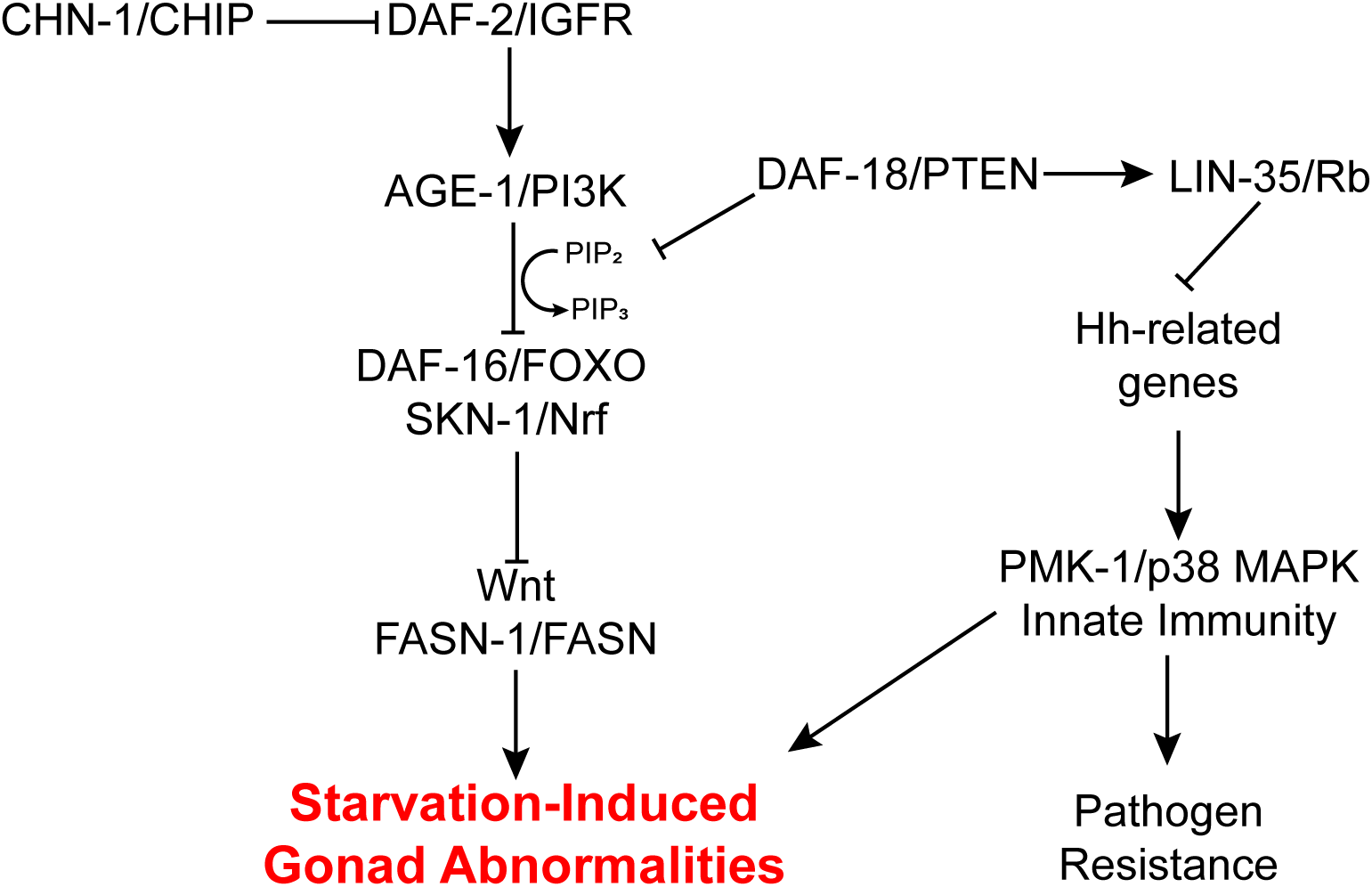
A model for how *daf-18/PTEN* and *lin-35/Rb* protect worms from developing L1 starvation-induced gonad abnormalities. Previous work has shown that reducing IIS suppresses starvation-induced abnormalities via *daf-16/FoxO* and *skn-1/Nrf*. We show that in addition to antagonizing IIS, *daf-18/PTEN* suppresses abnormalities via *lin-35/Rb*, which antagonizes Hh-related signaling. We also show that Hh-related signaling promotes abnormalities by activating *pmk-1/p38 MAPK*-mediated innate immunity. Consequently, in addition to causing developmental abnormalities that limit reproductive success, extended early life starvation promotes pathogen resistance later in life.

### *daf-18/PTEN* and *lin-35/Rb* function independently of insulin/IGF signaling to suppress starvation-induced germline tumors and other gonad abnormalities

PTEN is a potent tumor suppressor that is mutated in multiple advanced cancers at high frequency (Li *et al*. 1997; Steck *et al*. 1997), and partial loss of PTEN function, aberrant subcellular localization, and post-translational dysregulation have been linked to tumorigenesis and cancer progression (Lee *et al*. 2018). The sole worm PTEN homolog, *daf-18*, is required to repress cell division and other aspects of postembryonic development during L1 arrest (Fukuyama *et al*. 2006), and mutation of *daf-18* causes abnormal gonad development sterility following relatively brief L1 arrest (Wolf *et al*. 2014; Chen *et al*. 2022). Furthermore, *daf-18* RNAi during larval development after extended L1 arrest increases the frequency of starvation-induced gonad abnormalities (Shaul *et al*. 2023), suggesting that DAF-18/PTEN functions as a tumor suppressor beyond L1 arrest. *daf-16/FoxO* mutants have similar L1 arrest-related phenotypes to *daf-18*, but the phenotypes are less severe for *daf-16* (Fig. 1 and S1; Baugh and Sternberg 2006; Jordan *et al*. 2019; Chen *et al*. 2025). We show that *daf-18* suppression of starvation-induced gonad abnormalities depends in part on PI3K/IIS, but that *daf-18/PTEN* also functions independently of *age-1/PI3K* (Fig. 1). PTEN is a dual phosphatase, with lipid and protein-phosphatase activities (Myers *et al*. 1997, 1998). Protein-phosphatase activity of DAF-18/PTEN has not been demonstrated biochemically, but our results suggest such activity contributes to suppression of starvation-induced germ cell tumors and other gonad abnormalities. Alternatively, DAF-18 could dephosphorylate a lipid other than PIP3, or it could affect starvation-induced abnormalities non-enzymatically.

The retinoblastoma gene *RB1* was the first tumor suppressor identified (Berry *et al*. 2019), and it is mutated in many types of cancer (Burkhart and Sage 2008; Vélez-Cruz and Johnson 2017). We show that the sole worm RB homolog, *lin-35/Rb*, represses starvation-induced germ cell tumors (Fig. 1), revealing tumor-suppressor activity of LIN-35 for the first time. We recently found that *daf-18/PTEN* requires *lin-35/Rb* to promote starvation resistance during L1 arrest (Chen *et al*. 2025), leading us to hypothesize that *daf-18* acts through *lin-35* to suppress development of starvation-induced gonad abnormalities. We show that *lin-35* functions independently of *daf-2/InsR*, and that *lin-35* and *daf-18* do not have additive effects on starvation-induced abnormalities (Fig. 1). Furthermore, lack of additivity was observed with RNAi, indicating gene function during recovery from starvation. These results suggest that in addition to functioning in a linear pathway to promote starvation resistance (Chen *et al*. 2025), *daf-18* and *lin-35* function in a common pathway during larval development to suppress starvation-induced tumors.

### *daf-18/PTEN* and *lin-35/Rb* antagonize Hedgehog-related signaling, which cell-nonautonomously promotes starvation-induced abnormalities

Genes involved in Hedgehog signaling in *Drosophila* and mammals are conserved in *C. elegans*, but they are expanded into families of homologs with divergent structures. Among the *C. elegans* Hh-related genes, there are 60 putative ligands, only 10 of which possess the conserved C-terminal auto-processing domain, and 26 putative receptors, 24 of which are classified as “patched-related” (*ptr*) genes (Aspöck *et al*. 1999). *C. elegans* lacks a Smo homolog, and there is no evidence that the Gli homolog TRA-1 acts as a transcriptional effector for signaling, suggesting that *C. elegans* Hh-related signaling is non-canonical (Zugasti *et al*. 2005). The phenotypes reported for Hh-related signaling in *C. elegans* are not particularly reminiscent of the roles of Hh signaling in patterning, cell proliferation, and differentiation seen in *Drosophila* and mammals (Bürglin and Kuwabara 2006; Zhang and Beachy 2023). However, we show that four Hh-related genes, *ptr-23*, *wrt-1*, *wrt-10*, and *tra-1*, promote starvation-induced germ cell tumors and other gonad abnormalities (Fig. 2 and S4), suggesting conserved proto-oncogene function. In addition, we show that *ptr-23*, *wrt-1,* and *wrt-10* are epistatic to *daf-18/PTEN* and *lin-35/Rb* (Fig. 2), suggesting that DAF-18 and LIN-35 antagonize Hh-related signaling. Published transcriptome analysis of L3 larvae suggests that Hh-related genes are not differentially expressed in *lin-35* null mutants that had been continuously fed (Latorre *et al*. 2015), but such regulation may occur only in larvae that experienced L1 starvation. However, reporter gene analysis suggests that L1 arrest, *daf-18*, and *lin-35* do not have appreciable effects on transcription of *ptr-23, wrt-1*, or *wrt-10* (Fig. S3). Nonetheless, it is possible that multi-copy transcriptional reporters lack the sensitivity to capture relevant changes in Hh-related gene expression. It is unclear how LIN-35 affects Hh-related gene activity, but these observations suggest that LIN-35 regulates *ptr-23*, *wrt-1*, and *wrt-10* indirectly or post-transcriptionally. Notably, there is evidence in mammals that loss of *RB1* promotes aberrant cilia formation leading to hypersensitivity to Hh signaling (Cochrane *et al*. 2020), but it is unclear if this relevant to our work given the unciliated hypodermis as the apparent major site of action for the Patched receptor-related gene *ptr-23* (Fig. 3; see below).

Reporter gene analysis largely corroborated by published scRNA-seq results suggest that *ptr-23*, *wrt-1*, and *wrt-10* are prominently expressed in the hypodermis as well as the intestine and other relatively minor sites (Fig. 3). Germline expression was not observed, but the multi-copy transgenic arrays analyzed are likely silenced in the germline. Nonetheless, a pair of complementary, RNAi-based approaches suggest that all three genes function in the soma, and not the germline, to promote starvation-induced abnormalities (Fig. 3). All three Hh-related genes were required in the hypodermis to promote gonad abnormalities, and *wrt-1* was also required in the intestine. In addition, tissue-specific transgenic rescue of *ptr-23*, *wrt-1*, and *wrt-10* mutants suggests that over-expression of each gene in the hypodermis or intestine is sufficient to promote starvation-induced abnormalities (Fig. 3). Tissue-specific RNAi and transgenic rescue results are consistent with each other. These results are also consistent with the observed expression patterns, though other sites of expression were not tested for function (*e.g.*, ciliated neurons). Hypodermal function of Hh-related genes suggests the Hh-related signaling functions cell-nonautonomously to promote germ cell proliferation (Jordan *et al*. 2019). We do not know if PTR-23 functions as a receptor for WRT-1 or WRT-10, or if their activity is transduced through TRA-1/GLI as a transcriptional effector. In any case, cell-nonautonomous function suggests that PTR-23, WRT-1, and WRT-10 activity in the hypodermis, or possibly intestine, affects at least one other molecule that directly affects the somatic gonad or germ cells to influence their proliferation following extended L1 arrest.

Phenotypic and expression analysis suggests that *ptr-23*, *wrt-1*, and *wrt-10* have much in common, but *wrt-1* stands out in a few ways. *wrt-1* was unique in being necessary and sufficient in both the intestine and hypodermis to promote starvation-induced abnormalities (Fig. 3). *wrt-1* RNAi during development following extended starvation also affected more genes than *ptr-23* or *wrt-10* (or *tra-1*) (Fig. 4). In addition, mutation of *wrt-1* led to the most dramatic decrease in survival on *P. aeruginosa* PA14, which was significantly lower than wild type even in the absence of prior starvation (Fig. 5, see below). While *wrt-1* and *wrt-10* both possess a similar *wrt* N-terminal signaling domain, *wrt-1* possesses the conserved C-terminal auto-processing domain, which is absent in the majority of Hh-related putative ligands (Bürglin 1996, 2008). It is possible that the autocatalytic domain impacts post-translational regulation, secretion, and signaling function, accounting for the more severe effects of loss of *wrt-1*.

### Hedgehog-related signaling mediates activation of innate immunity by early life starvation, promoting developmental abnormalities but increasing pathogen resistance

RNA-seq analysis of adults that had been subjected to extended L1 arrest and then cultured with RNAi targeting *ptr-23*, *wrt-1*, *wrt-10*, or *tra-1* affected a relatively small number of largely overlapping genes (Fig. 4). Overlapping effects on gene expression suggest these Hh-related genes do in fact function in a common pathway. Furthermore, genes associated with innate immunity were significantly enriched among genes downregulated by disruption of Hh-related signaling. RNAi of such immunity-associated genes as well as the immunity regulator *pmk-1/p38 MAPK* reduced the frequency of starvation-induced abnormalities, even with hyperactivation of Hh-related signaling (Fig. 4), supporting the conclusion that *ptr-23*, *wrt-1*, and *wrt-10* promote transcription of immunity genes. Furthermore, relatively brief (1 d) or extended (8 d) L1 arrest caused upregulation of an immunity marker (*sysm-1p::GFP*) and increased pathogen resistance (Fig. 5 and S5), suggesting that the immunity-associated genes upregulated by Hh-related signaling actually function to promote immunity in this context. Connections between *C. elegans* Hh-related signaling and innate immunity have been documented (Zárate-Potes *et al*. 2022; Wang *et al*. 2023; VanDerMolen *et al*. 2025). Interestingly, both activation and repression of innate immunity have been observed in different contexts and for different Hh-related genes. Our results suggest that Hh-related signaling, the innate immunity pathway, and starvation-induced gonad abnormalities are causally related, but how an elevated immunity response following extended L1 arrest causes germ cell tumors and other gonad abnormalities is mysterious. Furthermore, the mechanism by which innate immunity is induced during recovery from starvation remains unclear. Our results suggest that the non-pathogenic *E. coli* used to feed the worms induces *sysm-1p::GFP* expression with or without L1 starvation, with a stronger induction in larvae that experienced starvation, and the strongest induction in larvae that experienced extended (8 d) starvation (Fig. 5). We suggest that previously starved larvae may have a stronger antigenic response to this commensal food source, rendering larvae subjected to L1 arrest relatively resistant to pathogenic bacteria during late larval development.

L1 arrest enables *C. elegans* larvae to endure long periods of starvation, which is likely ecologically relevant. However, extended L1 arrest has a variety of consequences that are seemingly pathological or maladaptive, including delayed development, reduced adult size, reduced fecundity, reduced progeny size and quality, and germ cell tumors and other gonad abnormalities in adults (Jobson *et al*. 2015; Jordan *et al*. 2019; Olmedo *et al*. 2020). Furthermore, animals that develop gonad abnormalities display the largest reduction in reproductive success (Jordan *et al*. 2019). In contrast, we report here that extended L1 arrest increases resistance to bacterial pathogens (Fig. 5 and S5), which does not seem pathological and could be adaptive. Increased pathogen resistance could increase fitness following starvation in environments where optimal food sources may not be available. Nonetheless, this potential evolutionary benefit is apparently offset by decanalization of development compromising adult phenotype and reproductive success.

## MATERIALS AND METHODS

### C. elegans strains

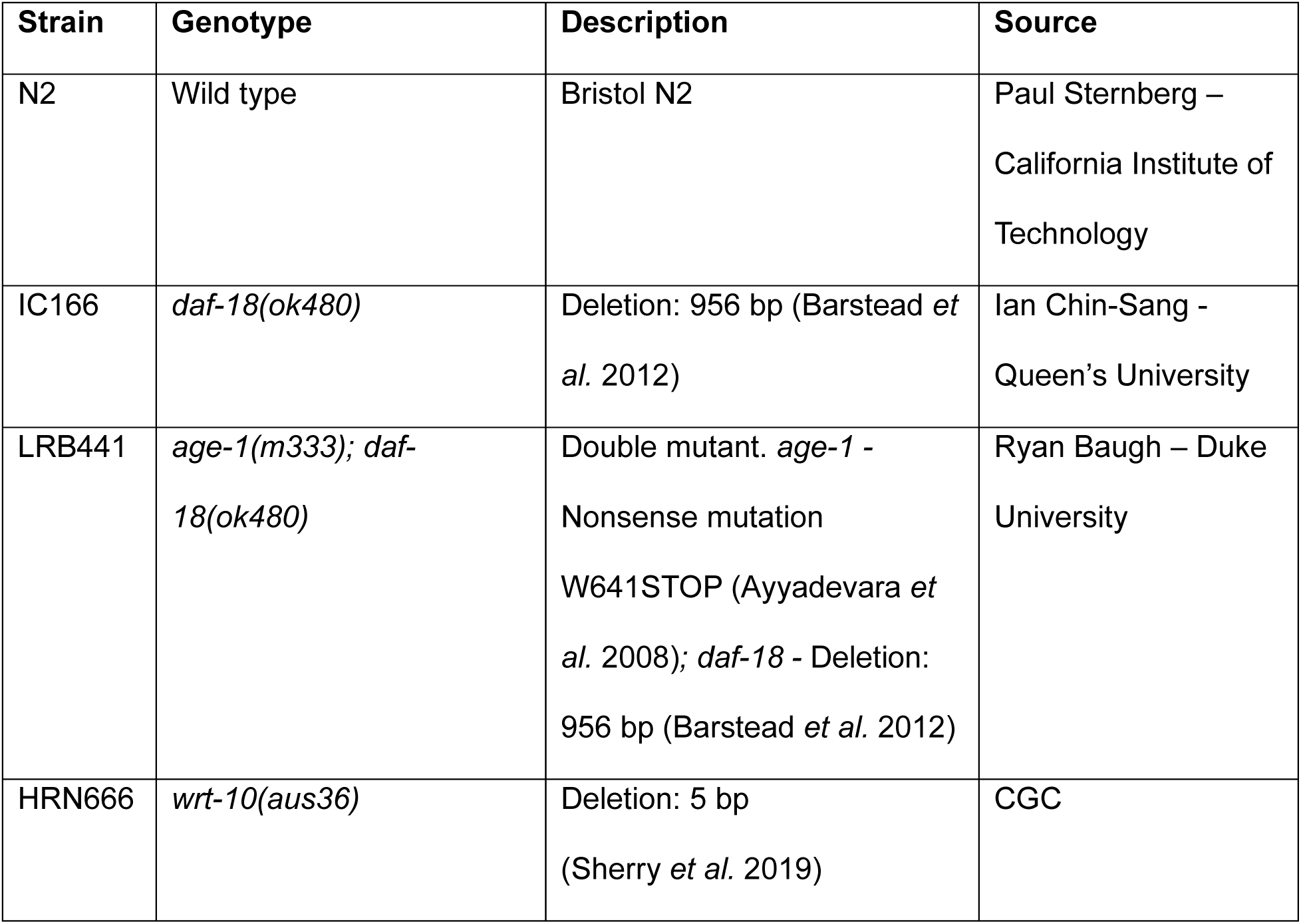
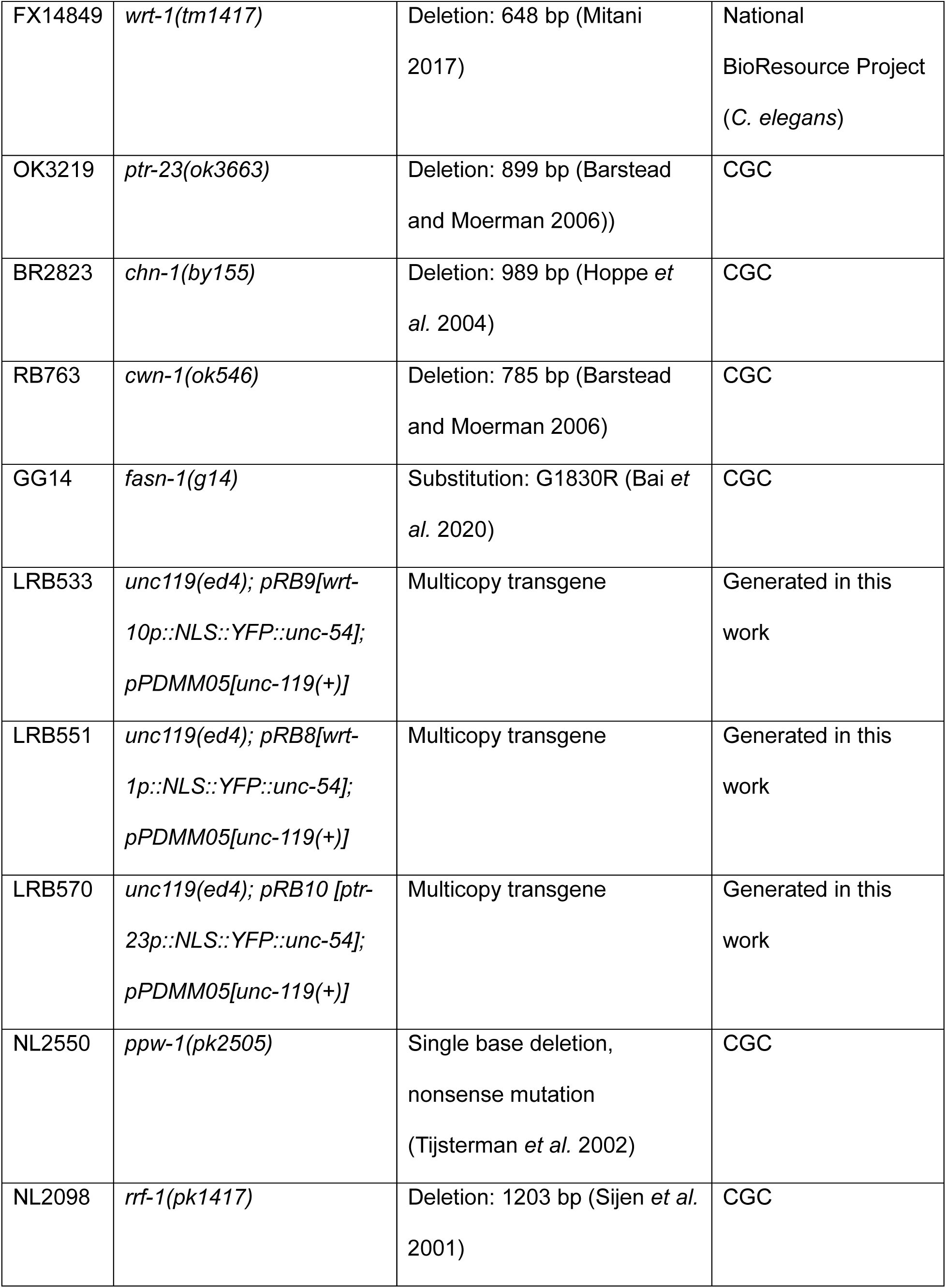
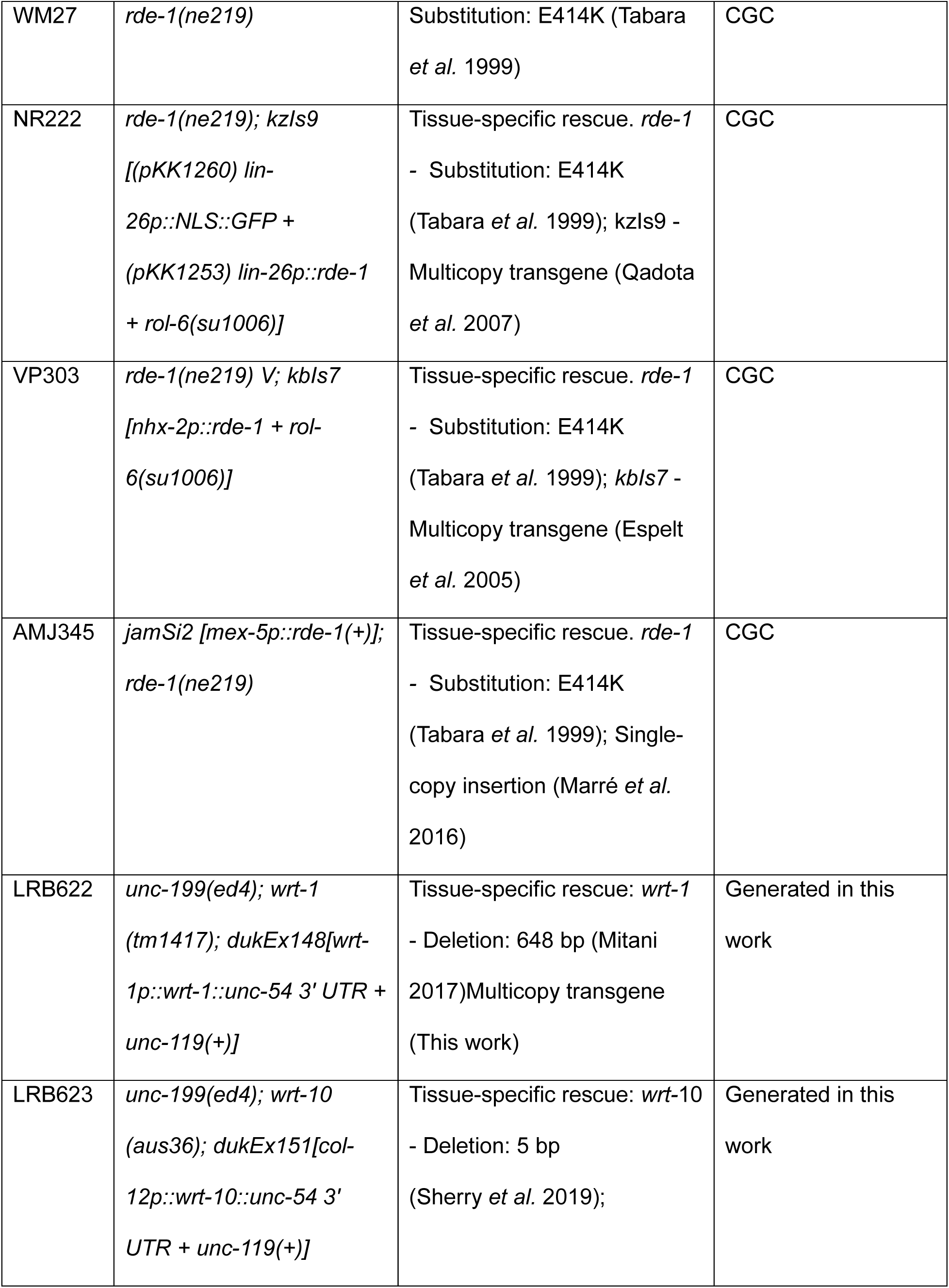
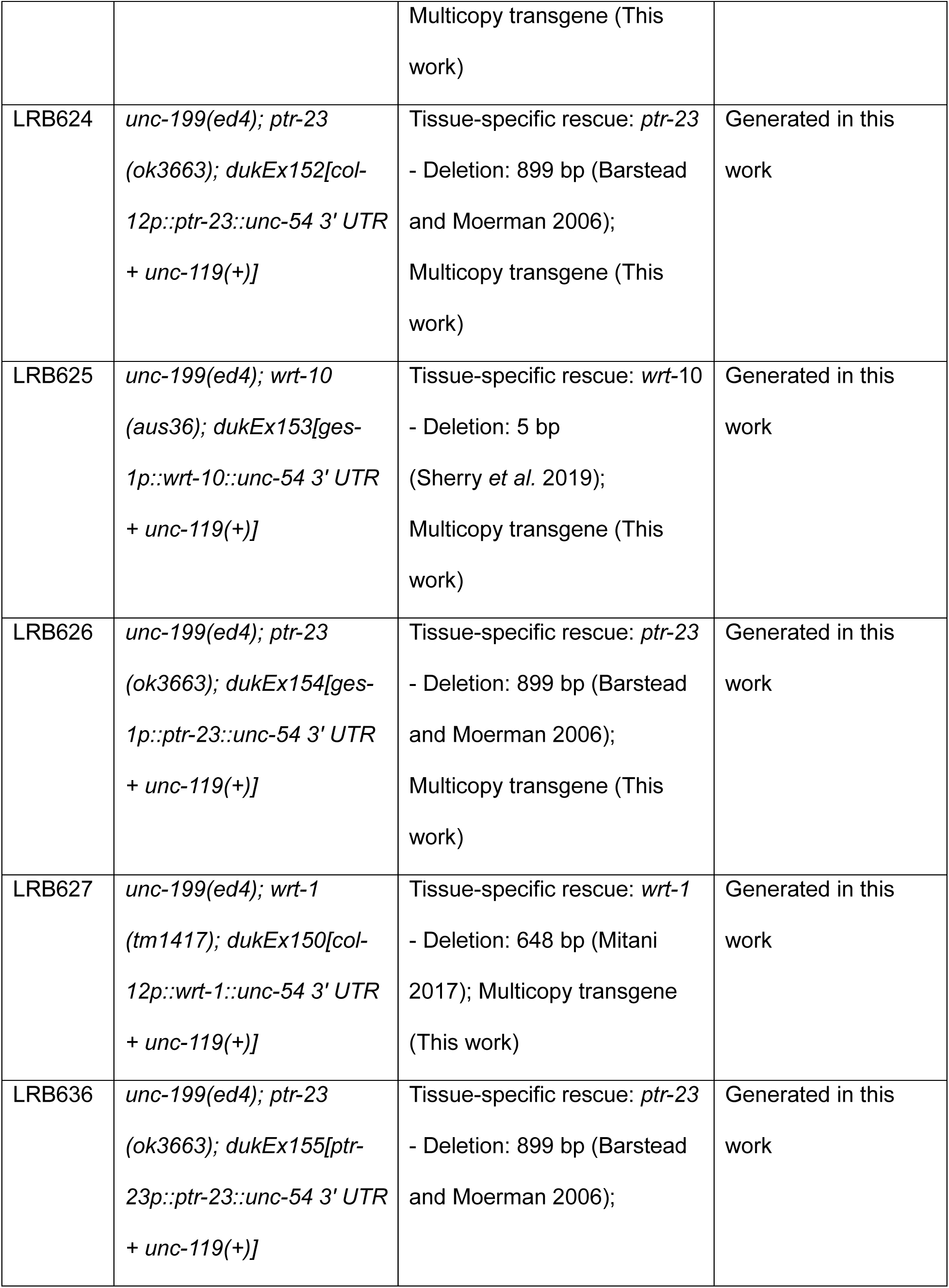
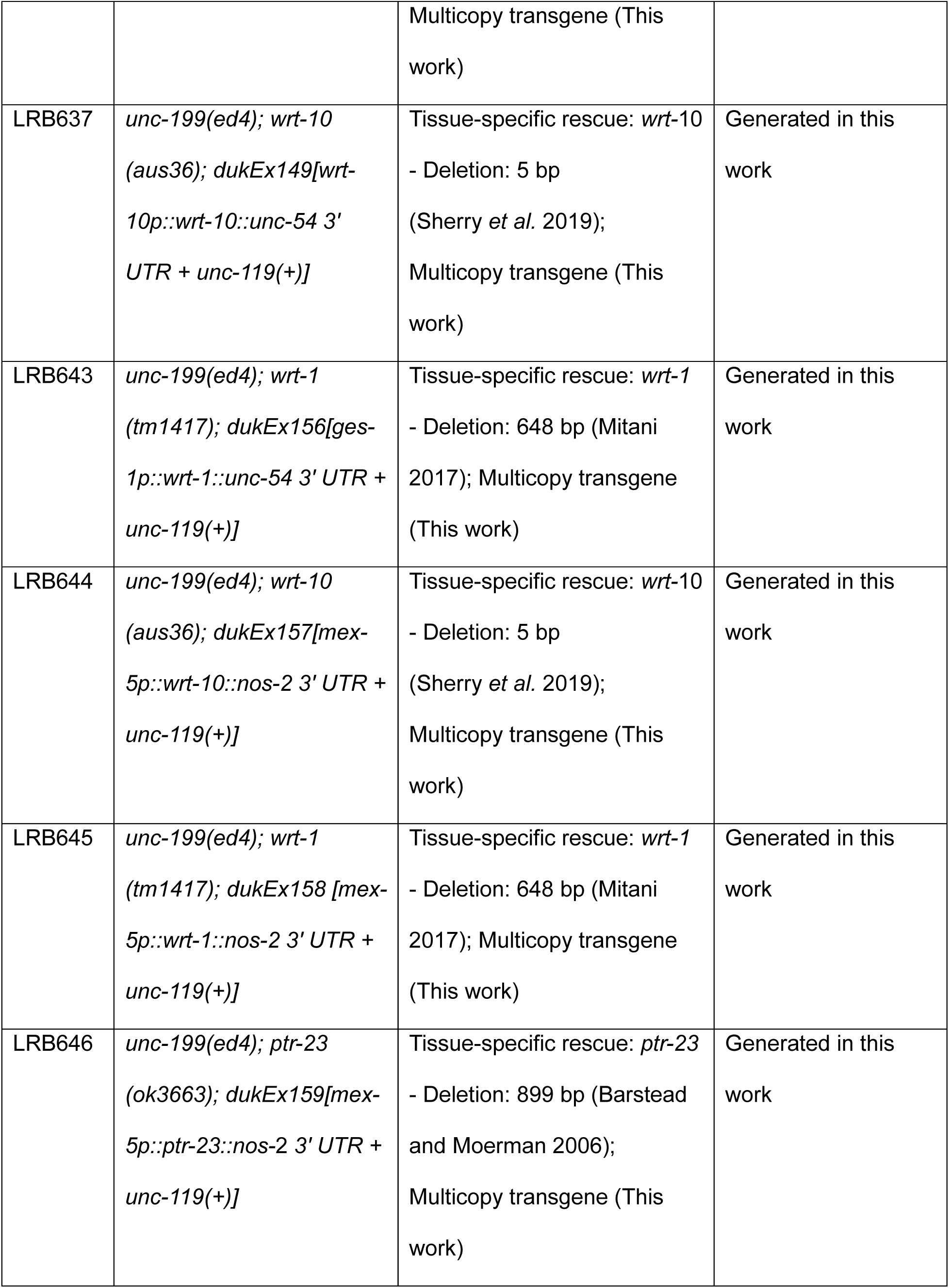
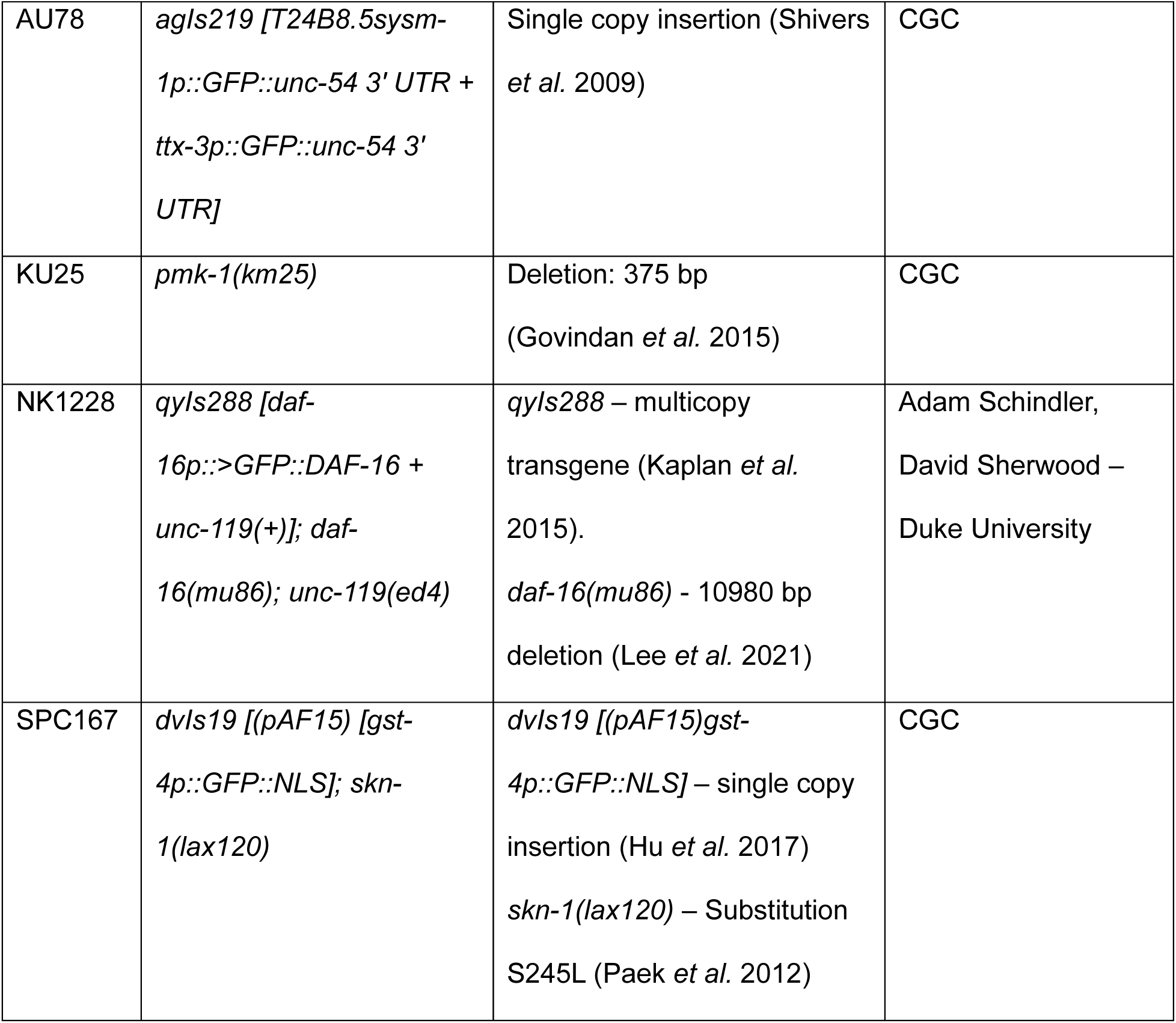

### Bacterial strains

| Species | Strain | Source |
| --- | --- | --- |
| *Escherichia coli* | OP50 | CGC |
| *Pseudomonas aeruginosa* | PA14 | Joel Myer – Duke University |
| *Salmonella enterica* | SL1344 | Joel Myer – Duke University |
| *Escherichia coli* | HT115 – L4440 empty vector | Colleen Murphy – Princeton University |
| *Escherichia coli* | HT115 – *daf-18* RNAi | Open Biosystems – David Sherwood - Duke University |
| *Escherichia coli* | HT115 – *lin-35* RNAi | Open Biosystems – David Sherwood - Duke University |
| *Escherichia coli* | HT115 – *daf-2* RNAi | Colleen Murphy – Princeton University |
| *Escherichia coli* | HT115 – *ptr-23* RNAi | Open Biosystems - David Sherwood - Duke University |
| *Escherichia coli* | HT115 – *wrt-1* RNAi | Open Biosystems - David Sherwood - Duke University |
| *Escherichia coli* | HT115 – *wrt-10* RNAi | Open Biosystems - David Sherwood - Duke University |
| *Escherichia coli* | HT115 – *tra-1* RNAi | Open Biosystems - David Sherwood - Duke University |
| *Escherichia coli* | HT115 – *dod-24* RNAi | Open Biosystems - David Sherwood - Duke University |
| *Escherichia coli* | HT115 - F55G11.8 RNAi | Open Biosystems - David Sherwood - Duke University |
| *Escherichia coli* | HT115 - F56A4.2 RNAi | Open Biosystems - David Sherwood - Duke University |
| *Escherichia coli* | HT115 – *oac-6* RNAi | Open Biosystems - David Sherwood - Duke University |
| *Escherichia coli* | HT115 - Y47H10A.5 RNAi | Open Biosystems - David Sherwood - Duke University |
| *Escherichia coli* | HT115 - C18H9.6 RNAi | Open Biosystems - David<br>Sherwood - Duke University |
| *Escherichia coli* | HT115 – *ugt-44* RNAi | Ahringer Library – David<br>Sherwood – Duke University |
| *Escherichia coli* | HT115 – *irg-5* RNAi | Ahringer Library – David<br>Sherwood – Duke University |
| *Escherichia coli* | HT115 – *cpr-3* RNAi | Ahringer Library – David<br>Sherwood – Duke University |
| *Escherichia coli* | HT115 - K08D8.5 RNAi | Ahringer Library – David<br>Sherwood – Duke University |
| *Escherichia coli* | HT115 – *asp-14* RNAi | Ahringer Library – David<br>Sherwood – Duke University |
| *Escherichia coli* | HT115 – *clc-19* RNAi | Ahringer Library – David<br>Sherwood – Duke University |

### Worm maintenance

Standard culture methods were carried out at 20°C on Nematode Growth Media (NGM) agar plates seeded with *E. coli* OP50. For the gonad abnormality assay, reporter quantification, and pathogen resistance assay, plates were seeded with *E. coli* HT115 empty vector bacteria (see below) or other indicated RNAi bacteria. All strains were cultured for at least five generations in well-fed conditions at 20°C prior to use in any experiments. All mutants used were backcrossed to wild type N2 for at least 4 generations.

### Hypochlorite treatment and starvation culture preparation

For starvation cultures used for scoring gonad abnormalities, sterile, synchronized embryos were obtained by hypochlorite treatment of adults on the first day of egg laying. Eight adults on the first day of egg laying were transferred to a 10 cm NGM plate seeded with *E. coli* OP50. After 72 hours at 20°C, progeny (in their first day of egg laying) were washed from the plates with S-basal medium and hypochlorite-treated to obtain embryos. For all other assays, seven L4 larvae were transferred and grown for 96 hours prior to hypochlorite treatment (Stiernagle 2006). Unless otherwise noted (Fig. 1A), embryos were transferred to virgin S-basal (without ethanol and cholesterol) and maintained at 20°C on a tissue-culture roller drum at a density of 1 animal per µL to hatch and enter L1 arrest.

### RNA interference

*E. coli* HT115 bacteria carrying a plasmid expressing double-stranded RNA for the indicated gene (or the empty vector control L4440) were used in all treatments. All RNAi bacteria used are listed above. Glycerol stocks were stored at -80°C. Frozen stocks were streaked on Luria Bertani (LB) plates with carbenicillin (Carb; 100 mg/ml) and tetracycline (Tet; 12.5 mg/ml) and grown overnight at 37°C. A single colony was picked into 1 mL LB + Carb (50 mg/ml) + Tet (12.5 mg/ml) and grown overnight at 37°C and 250 rpm. This culture was used to inoculate 5-20 mL of Terrific Broth (TB) with Carb (50 mg/mL) and grown overnight at 37°C and 300 rpm. Turbid cultures were centrifuged for 10 minutes at 4000 rpm and the supernatant was removed. Bacteria were resuspended in S-complete + 20% glycerol at a 4:1 ratio by mass. 100 µL aliquots were frozen and stored at -80°C. 15 µL from one of these aliquots was used to inoculate 6 cm NGM + Carb (50 mg/ml)+ IPTG (1 mM) plates and grown overnight at room temperature before use in experiments.

### Scoring gonad abnormalities

For extended starvation conditions, L1 larvae were typically arrested in virgin S-basal for 8 days after hypochlorite treatment, but in limited cases they were arrested for 4 days and/or 0.1% ethanol was added to S-basal (Fig 1A). Two types of control conditions were used, as indicated: 1) Embryos were allowed to hatch and arrest overnight for synchronization, approximately 18 hours after hypochlorite treatment (“1 day L1 arrest”), or 2) embryos were plated directly with food so they hatch and proceed directly to postembryonic development without entering L1 arrest (“Unstarved”). 150 arrested L1 larvae or unhatched embryos were transferred to 6 cm NGM + Carbenicillin + IPTG plates seeded with *E. coli* HT115 expressing an empty vector RNAi plasmid (L4440) or indicated RNAi treatment. Worms were grown to early adulthood (approximately 60 hours for animals starved for 18 hours, 68 hours for animals starved for 4 days, and approximately 72 hours for animals starved for 8 days or unhatched embryos) then washed with S-basal and anesthetized with levamisole (10 mM). Paralyzed adults were transferred to 4% noble agar pads on microscope slides and viewed at 200X total magnification using differential interference contrast/Nomarski (DIC) microscopy on a Zeiss AxioImager compound microscope. Fifty individuals were scored for the presence of proximal germ cell tumors, uterine masses, or other obvious gonad abnormalities (Fig. S1A, B (Falsztyn *et al*. 2025)). Proximal germ cell tumors present as large masses near the vulva/proximal gonad with visible germ cell nuclei (similar to the mitotic distal gonad), and uterine masses are characterized as large irregular tissues in the uterus without visible germ cell nuclei (differentiated) often resulting in extruded vulvae. For statistical analysis, Bartlett’s test was used to determine homogeneity of variance, and if non-significant, variance was pooled for subsequent analysis. Two-tailed, unpaired t-tests were used for pairwise comparisons.

### Plasmid design and cloning

Gibson assembly was used for making new plasmids. Q5 high-fidelity polymerase PCR (NEB M0491) was used to amplify plasmid fragments from plasmid DNA and promoter sequences from wild-type genomic DNA (see Table S6 for PCR primer sequences). DNA sequences for design were obtained from WormBase (Sternberg *et al*. 2024). Hedgehog-related gene coding sequences were obtained from genomic DNA, YFP reporter genes with the plasmid backbone were sub-cloned from pPD132.112 (Andy Fire Lab Vector Kit), and the pNL213, pAS10, and pGC550 plasmids, which include promoters and 3’ UTRs for intestine, hypodermis, and germline expression, respectively, were used as vectors. PCR fragments were gel purified (Zymo Research D4001). Purified DNA was used for Gibson assembly based on the kit manufacturer instructions, and the resulting products were used for bacterial transformation using competent cells from the same kit (NEB E5510S). Plasmid DNA was extracted from individual colonies (Zymo Research D4016), and plasmids were sequenced to confirm their structure.

### Plasmid microinjection

Microinjection was used to generate all multicopy transcriptional reporter lines and tissue-specific rescue lines used in this work. Worms carrying the *unc-119(ed4)* mutation were used for all injections. Young adults (not yet gravid) or late L4 larvae were picked onto unseeded (lacking bacterial food) NGM plates. A paintbrush and halocarbon oil were used to transfer worms onto a desiccated 2% noble agar pad on a 45 mm x 50 mm cover glass. Plasmids were injected into the distal gonad at a concentration of 100 ng/μl. An *unc-119* rescue plasmid (pPDMM051) was used as an injection marker at a concentration of 50 ng/μl. Injected animals were transferred to 2 cm NGM plates with seeded with *E. coli* OP50, and transformants were identified among progeny by *unc-119* rescue (wild type morphology and motility). Transformed progeny were singled to new plates, and stable lines were genotyped for validation by confirming GFP expression or PCR targeting the promoter-coding sequence fusion in the transgene.

### Compound microscopy imaging of reporter genes

Well-fed L4 larvae were transferred to 2 μL of 10 mM Levamisole on a 4% noble agar pad on a microscope slide and covered with a coverslip. Worms were imaged at 200x (AU78) or 400X (LRB533, LRB551, LRB570) total magnification using a Zeiss AxioImager compound microscope with an AxioCam 506 Mono camera. FIJI and Adobe Illustrator were used for miscellaneous editing, cropping, and stitching of images.

### RNA-seq sample collection

Starvation cultures were prepared as for scoring gonad abnormalities as described above. Following 8 days of starvation, 150 L1 larvae were plated onto empty vector, *ptr-23, tra-1, wrt-1,* and *wrt-10* RNAi. After 68 hours, adults were washed with S-basal from plates then centrifuged at 3000 rpm for 1 minute. Worms were washed in 10 mL of S-basal three additional times, and a final volume of 100 μL was transferred to a 1.5 mL Eppendorf tube. Samples were frozen in liquid nitrogen and stored at -80°C.

### RNA extraction and library preparation

100 μL of acid-washed sand and 1 mL of TRIzol were added to frozen worm pellets, and the tubes were vortexed vigorously for 10 minutes to homogenize samples. 200 μL of phenol-chloroform was added, and the mixture was vigorously vortexed for an additional 3 minutes.

Tubes were centrifuged for 3 minutes at 21,000 rpm. A p200 micropipette was used to extract the aqueous layer, which was transferred to a new 1.5 mL Eppendorf tube and mixed with 500 μL of isopropanol. The mixture was incubated for 8 minutes at room temperature followed by 2 minutes on ice. Tubes were subsequently centrifuged for 10 minutes at 21,000 rpm. RNA pellets were washed with 75% ethanol twice, after which they were allowed to air dry prior to resuspension in 10 μL of water. RNA quality and quantity were analyzed on NanoDrop and Qubit instruments, respectively. Libraries were prepared for sequencing using the NEBNext Ultra II RNA Library Prep Kit for Illumina (New England Biolabs #E7775) starting with 1 μg of total RNA per sample as input and seven cycles of PCR. Individually barcoded libraries were pooled and sequenced on the NovaSeq 6000 S-Prime flow cell to obtain 50 bp paired-end reads.

### RNA-seq analysis

Bowtie was used for mapping reads to the WS273 *C. elegans* genome and HTSeq was used for counting reads and generating count tables using a Linux command line on Ubuntu. EdgeR was used to estimate dispersion and perform an exact test for differential expression analysis on R Windows. A false discovery rate < 0.1 was used as a significance threshold for pairwise differential gene expression. A generalized linear model with a p-value cutoff < 0.05 was used for identifying differentially expressed genes across all treatments for clustering by log_2_ fold-change compared to the empty vector control. WormCat (Holdorf *et al*. 2020) was used for gene set enrichment analysis.

### Pathogen resistance

Fast-killing NGM plates (Tan *et al*. 1999) were seeded with 15 μL of *P. aeruginosa* PA14 or *S. enterica* SL1344 bacteria, which was spread to cover the entire plate surface, and incubated overnight at 37°C. Synchronized populations of unstarved L1s and L1s starved for 1 or 8 days were prepared as described above. 500 L1 larvae/embryos were plated onto *E. coli* HT115 bacteria and cultured for 56 hours for embryos, 48 hours for larvae starved for 1 day, and 56 hours for larvae starved for 8 days at 20°C. Fifty L4 larvae were picked onto pathogen plates. Survival was scored by observing movement. If worms were unresponsive to gentle prodding with a pick, they were removed from the plate and scored as dead. Worms that died from other causes, such as crawling onto the side of the plates, were censored. Survival was scored every 2 hours for the first 6 hours, 18 hours later (24 hours total), 6 hours later (30 hours total), 18 hours later, 6 hours later, and so on until no live worms remained. Oasis 2 (Han *et al*. 2016) was used for statistical analysis and the log-rank test was used for pairwise comparisons between strains.

### Automated imaging and quantitive analysis of reporter gene expression

Synchronized populations of unstarved L1s and L1s starved for 1 or 8 days were prepared as described above. 500 L1 larvae/embryos were plated onto empty vector *E. coli* HT115, or the indicated RNAi treatment, and cultured for 56 hours for embryos, 48 hours for L1 larvae starved for 1 day, and 56 hours for L1 larvae starved for 8 days, or the indicated duration, at 20°C. Worms were washed from plates with virgin S-basal and briefly centrifuged (15 seconds at 3000 rpm). Worms were resuspended in 390 µL of 50 μM sodium azide and transferred to a 96-well plate. Plates were imaged on an ImageXpress® Nano automated imager at 100X total magnification. Sites for the same well were stitched and objects resulting from automatic segmentation were analyzed to measure average fluorescent intensity. Objects were filtered by size (5 µm < width < 40 µm and 10000 < pixels < 75000 ), and background intensity was subtracted. Manual screening of objects also removed any instances of multiple animals or other debris. A linear mixed-effects model (ex: lme(intensity∼strain random = ∼1|replicate, data = imager) was used to fit average intensity per individual as the response variable, strain as the fixed effect, and replicate as the random effect for pairwise comparisons.

### Starvation survival

Starvation cultures were prepared as described above. After 1 day, 100 μL of the starvation culture was plated onto a 2 cm NGM plate seeded with *E. coli* OP50, and the number of plated L1 larvae was counted. After 48 hours at 20°C, the number of live larvae (exhibited growth or were still actively moving on plate surface) was counted, and the proportion of survivors was determined by dividing the number plated by the number of survivors. Survival was scored every 24 hours until there were no survivors. Quasi-binomial logistic regression, with the frequency of survivors as the response variable and the duration of starvation as the explanatory variable, was used for curve fitting and to estimate half-lives for each biological replicate. Bartlett’s test was used to determine if variance in replicate half-lives was homogeneous across genotypes, and if it was then variance was pooled across genotypes for subsequent analysis. Two-tailed, unpaired t-tests on replicate half-lives were used for pairwise comparisons between genotypes.

### Data availability

The authors affirm that all data necessary for confirming the conclusions of the article are present within the article, figures, and tables. Table S1 contains summary statistics for all gonad abnormality assays. Table S2 contains summary statistics and replicate-level data for starvation survival assays presented in Fig. S2. Table S3 contains all RNA-Seq data and analysis results presented in Fig. 4. Raw RNA-seq data is available from NCBI GEO (accession number GSE290428). Table S4 contains summary statistics and replicate-level data for all pathogen resistance assays. Strains are available upon request.

## ACKNOWLEDGEMENTS

We would like to thank David Sherwood’s lab for generously providing bacterial strains for RNAi. *C. elegans* strains were provided by the CGC, which is funded by NIH Office of Research Infrastructure Programs (P40 OD010440). We would also like to thank WormBase.

## Supporting Information Captions

**Figure S1.**
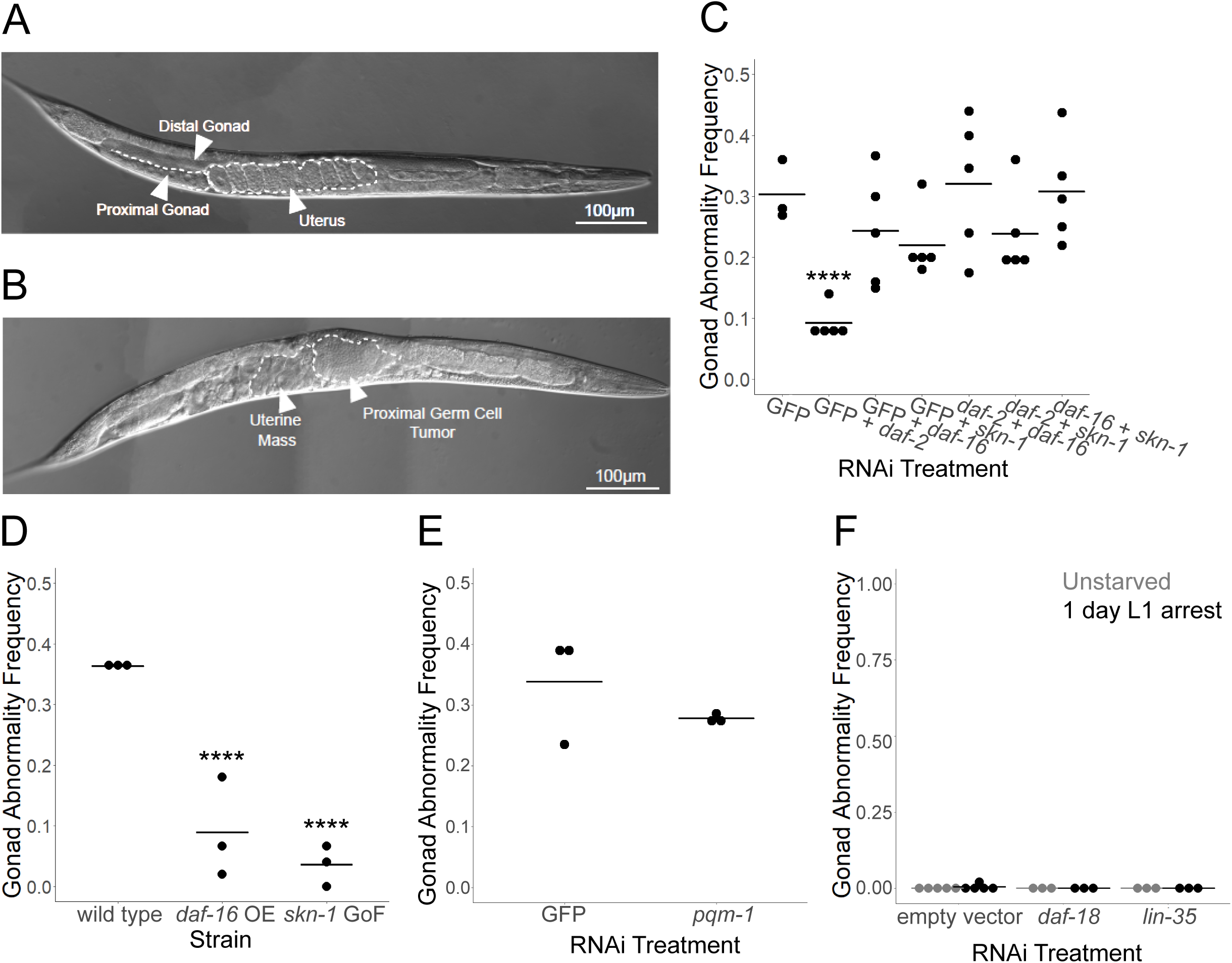
IIS effector genes do not account for the relatively strong effect of *daf-18/PTEN* on starvation-induced gonad abnormalities (related to Fig. 1). A) Representative image of a healthy adult without obvious gonad abnormalities after expended L1 arrest. B) Representative image of an adult with the most common gonad abnormalities observed after extended L1 arrest, including a differentiated uterine mass, a proximal germ cell tumor. A,B) Images of wild-type adults following extended L1 arrest (8 days) were taken at 400x total magnification 72 hr after plating with food at 20°C. Images were taken with differential interference contrast (DIC). FIJI and Adobe Illustrator were used for adjusting brightness/contrast and stitching images. Relevant tissues, organs, and abnormalities are outlined or indicated with arrowheads and labels. C, E) Wild type larvae were starved during L1 arrest for 8 days then recovered with RNAi bacteria targeting the indicated genes and scored for gonad abnormalities. D) Wild type, *daf-16(mu86);* a *daf-16* over-expression (OE) strain (*qyIs288 [daf-16p::GFP::DAF-16]*), and a *skn-1* gain-of-function (GoF) strain (*skn-1(lax120)*) were starved for 8 days then recovered on empty vector RNAi bacteria and scored for gonad abnormalities. F) Wild type larvae were starved for the indicated duration then recovered on the indicated RNAi bacteria and scored for gonad abnormalities. C-F) Each dot represents a biological replicate including ∼50 individuals (see Table S1 for summary statistics). Horizontal bars represent the mean across replicates. Asterisks indicate statistically significant differences between RNAi treatments and GFP (C), between mutants and wild type (B) (no statistical significance in E or F). **** P < 0.0001; unpaired, two-tailed t-test.

**Figure S2.**
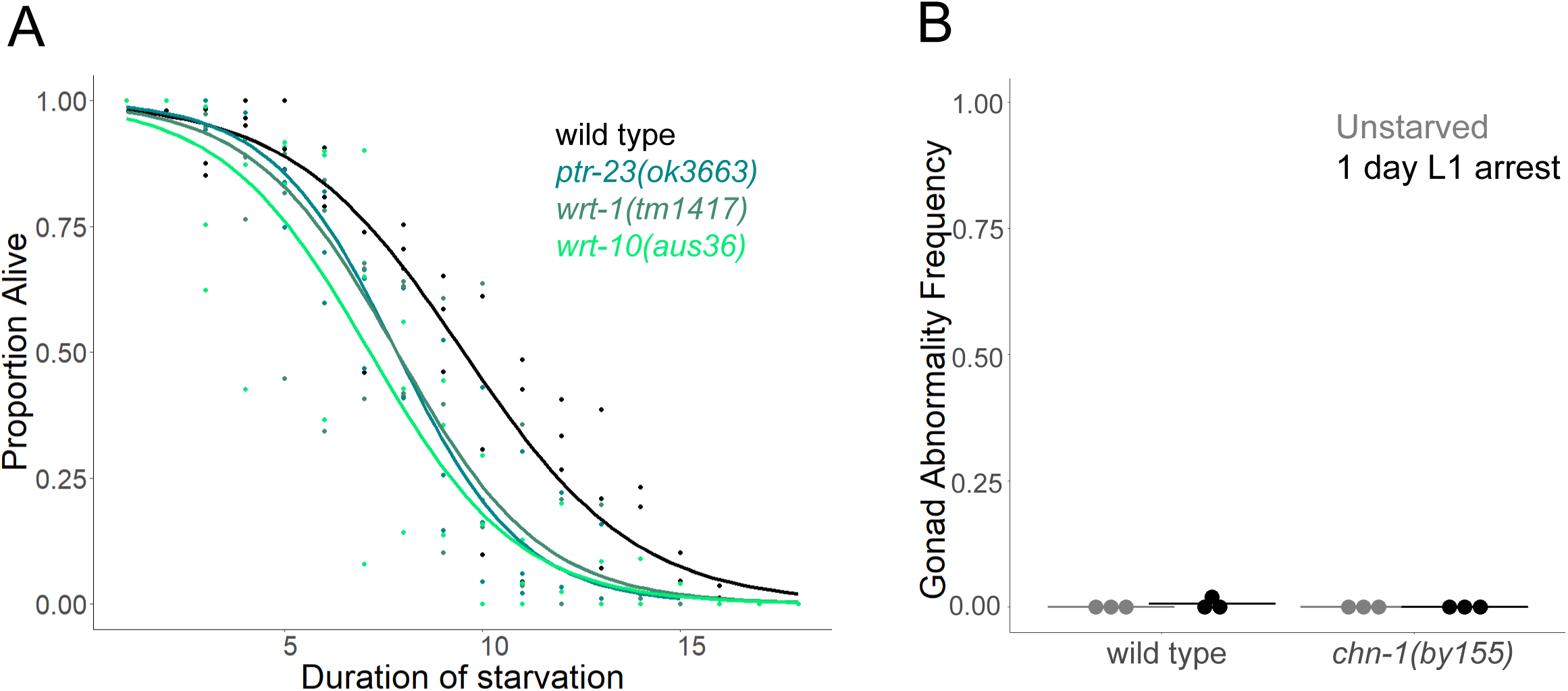
*ptr-23*, *wrt-1*, and *wrt-10* mutants do not have increased L1 starvation survival, and the *chn-1* mutant does not enhance abnormalities without extended L1 arrest (related to Fig. 2). A) L1 starvation survival was scored for the indicated genotypes in three biological replicates. ∼100 individuals (median = 92, range = 51-186) were scored per time point in each genotype. Curves were fit to each genotype with logistic regression and were used to calculate half-lives per replicate for each genotype, and an unpaired two-tailed t-test was used for pairwise comparisons of half-lives between each mutant and wild type. B) Wild type and *chn-1(by155)* larvae were starved for the indicated duration then recovered on empty vector RNAi and scored for gonad abnormalities. Each dot represents a biological replicate with ∼50 individuals (see Table S1 for summary statistics). Horizontal bars represent the mean across replicates. An unpaired, two-tailed t-test was used for pairwise comparisons between genotypes with the same duration of starvation. A-B) “n.s.” = not significant.

**Figure S3.**
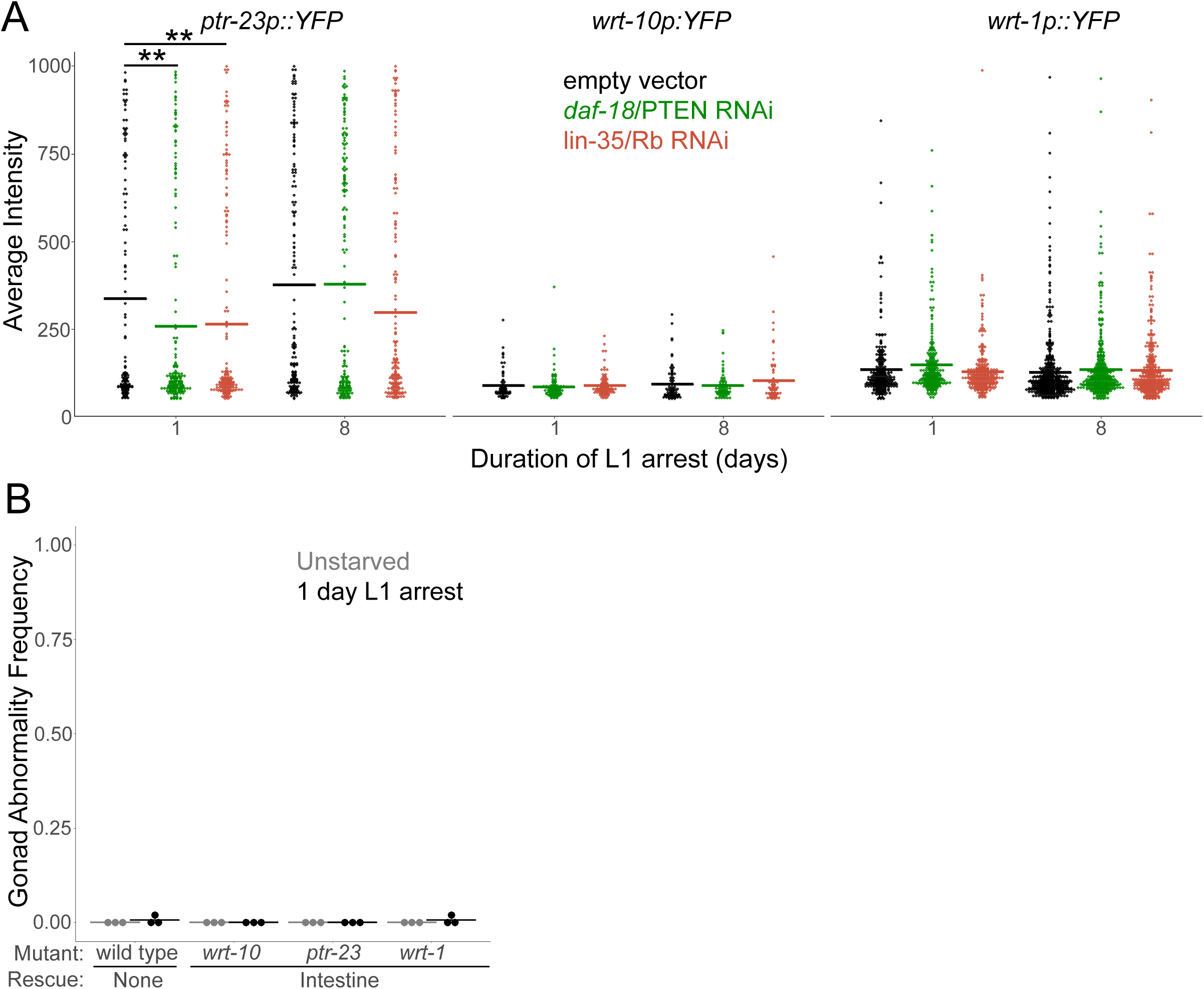
Transcription of *wrt-10*, *ptr-23*, and *wrt-1* is largely unaffected by L1 arrest, *daf-18/PTEN*, or *lin-35/Rb*, and overexpression of *wrt-10*, *ptr-23*, and *wrt-1* does not cause gonad abnormalities without extended L1 arrest (related to Fig. 3). A) L1 larvae with *ptr-23p::YFP*, *wrt-1p::YFP*, or *wrt-10p::YFP* reporter gene were starved for the indicated duration, recovered on plates with empty vector, *daf-18* RNAi, and *lin-35* RNAi bacteria for 48 hr, and whole-worm reporter gene expression was quantified by image analysis. To ensure matching stages, worms starved for 8 d were allowed an additional 8 hr to account for developmental delay following extended starvation. Individual points represent the average, background-corrected pixel intensity for a single worm. Horizontal bars represent the mean intensity across three biological replicates. Asterisks and bars indicate significance. A linear mixed-effect model was fit to the data with background-corrected average intensity as the response variable, RNAi treatment as the fixed effect, and replicate as the random effect. * P < 0.05, *** P < 0.001, **** P < 0.0001. C) Wild type and *ptr-23(ok3663), wrt-1(tm1417),* and *wrt-10(au36)* mutants rescued with a multicopy transgene with tissue-specific expression in the intestine (*ges-1p*) were starved for the indicated duration then recovered on empty vector RNAi bacteria and scored for gonad abnormalities. Each dot represents a biological replicate with ∼50 individuals (see Table S1 for summary statistics). Horizontal bars represent the mean across replicates. “n.s.” = not significant; unpaired, two-tailed t-test between each mutant and wild type with the same duration of starvation.

**Figure S4.**
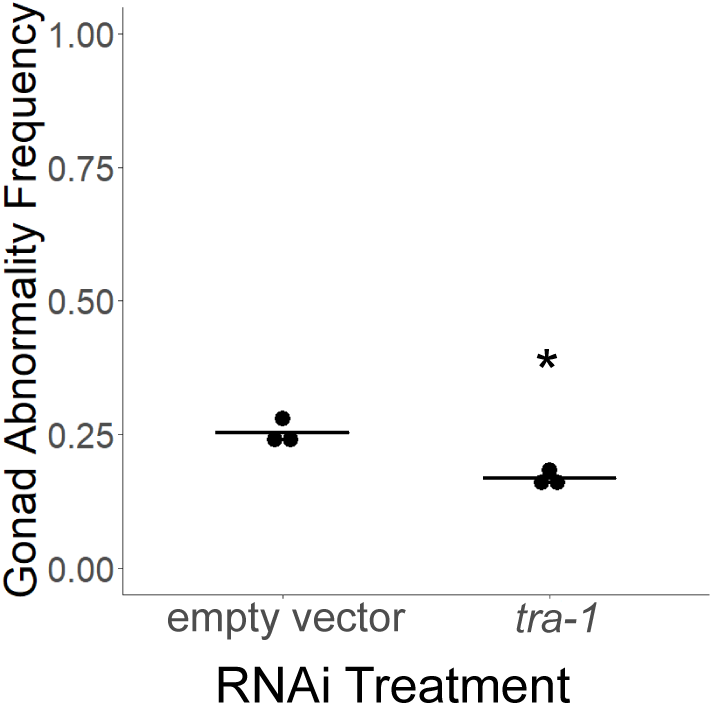
*tra-1/GLI* knockdown suppresses starvation-induced gonad abnormalities (related to Fig. 4). Wild type L1 larvae were starved for 8 days then recovered on empty vector RNAi bacteria or *tra-1* RNAi and scored for gonad abnormalities on the first day of egg laying. Each dot represents a biological replicate with ∼50 individuals (see Table S1 for summary statistics). Horizontal bars represent the mean across replicates. * P < 0.05; unpaired, two-tailed t-test comparing *tra-1* RNAi to empty vector.

**Figure S5.**
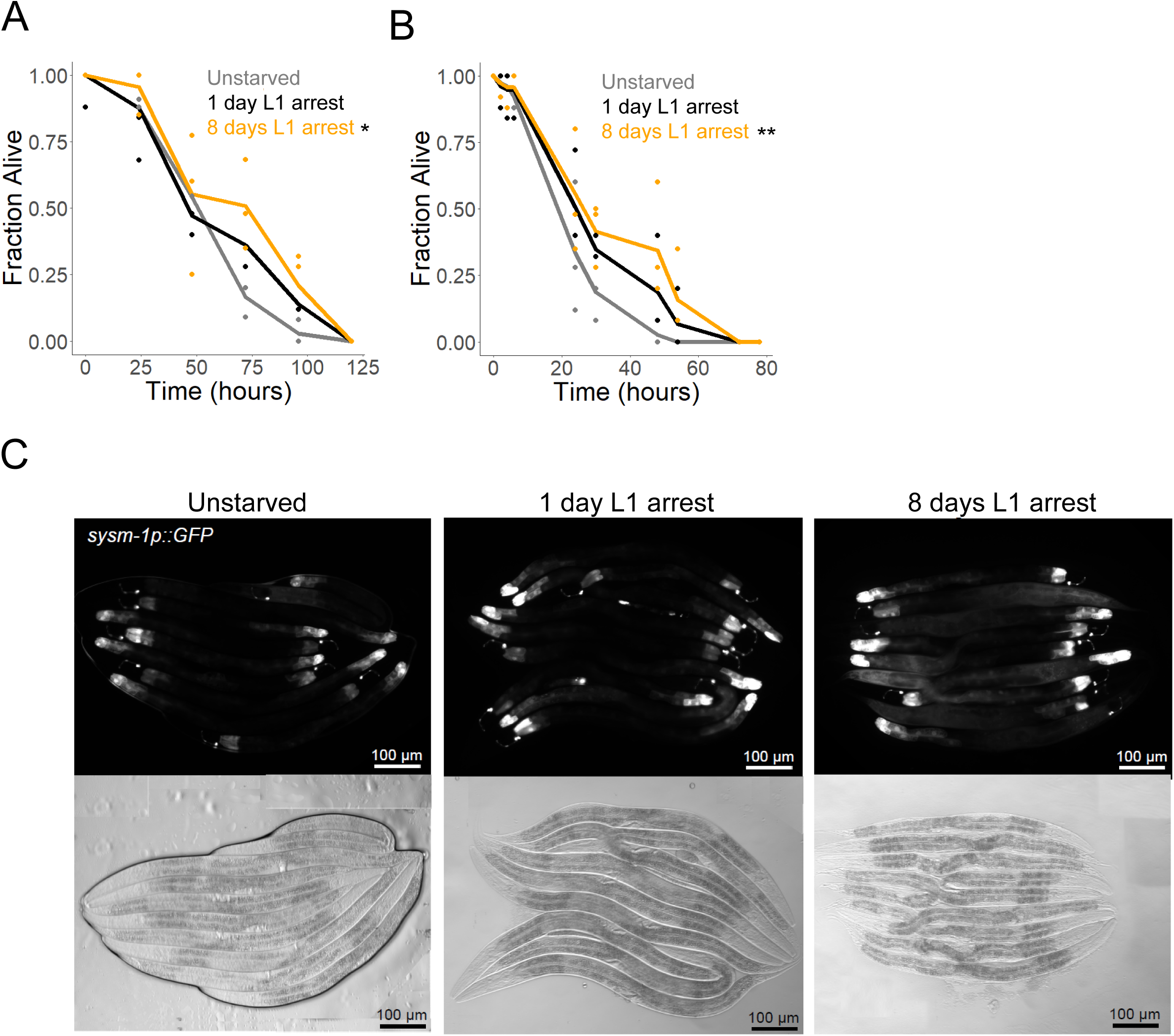
Starvation during L1 arrest activates innate immunity later in life (related to Fig. 5). A, (B) Wild-type L1 larvae were starved for the indicated duration, recovered on plates with *E. Coli* HT115 bacteria, L4 larvae were transferred to *P. aeruginosa* PA14 “slow-killing” plates (A) or *S. enterica* plates (B), and survival was assayed. Three biological replicates were performed with ∼50 individuals each (see Table S1 for summary statistics). Mean survival at each timepoint is plotted as a line with survival of each replicate at each timepoint included as dots. Asterisks indicate significance between the indicated starvation durations and unstarved. The log-rank test was used for pairwise comparisons; * P < 0.05, ** P < 0.01. C) Images of *sysm-1p::GFP* expression with a total magnification of 200x were taken of a random population of L4 larvae starved for the indicated duration (as in Fig. 5B). To ensure matching stages, worms starved for 0 hours were allowed an additional 8 hr to account for embryogenesis, and worms starved for 8 d were also allowed an additional 8 hr to account for developmental delay following extended starvation. Corresponding DIC images are also provided.

## REFERENCES

Aspöck G., H. Kagoshima, G. Niklaus, and T. R. Bürglin, 1999 Caenorhabditis elegans has scores of hedgehog-related genes: Sequence and expression analysis. Genome Res. 9: 909–923. 10.1101/gr.9.10.909

Ayyadevara S., R. Alla, J. J. Thaden, and R. J. Shmookler Reis, 2008 Remarkable longevity and stress resistance of nematode PI3K-null mutants. Aging Cell 7: 13–22. 10.1111/j.1474-9726.2007.00348.x

Bai X., D. Woodbury, and A. Golden, 2020 The fasn-1(g14ts) allele is a Gly1830Arg missense mutation in C. elegans FASN-1. microPublication Biol. 2020. 10.17912/micropub.biology.000244

Barstead R. J., and D. G. Moerman, 2006 C. elegans deletion mutant screening. Methods Mol. Biol. 351: 51–58. 10.1385/1-59745-151-7:51

Barstead R., G. Moulder, B. Cobb, S. Frazee, D. Henthorn, et al., 2012 Large-scale screening for targeted knockouts in the caenorhabditis elegans genome. G3 Genes, Genomes, Genet. 2: 1415–1425. 10.1534/g3.112.003830

Baugh L. R., and P. W. Sternberg, 2006 DAF-16/FOXO Regulates Transcription of cki-1/Cip/Kip and Repression of lin-4 during C. elegans L1 Arrest. Curr. Biol. 16: 780–785. 10.1016/j.cub.2006.03.021

Baugh L. R., 2013 To grow or not to grow: Nutritional control of development during Caenorhabditis elegans L1 Arrest. Genetics 194: 539–555. 10.1534/genetics.113.150847

Baugh L. R., and P. J. Hu, 2020 Starvation responses throughout the caenorhabditis elegans life cycle.

Berry J. L., A. Polski, W. K. Cavenee, T. P. Dryja, A. Linn Murphree, et al., 2019 The RB1 story: Characterization and cloning of the first tumor suppressor gene. Genes (Basel). 10. 10.3390/genes10110879

Bharill P., S. Ayyadevara, R. Alla, and R. J. Shmookler Reis, 2013 Extreme depletion of PIP3 accompanies the increased life span and stress tolerance of PI3K-null C. elegans mutants. Front. Genet. 4: 1–11. 10.3389/fgene.2013.00034

Bürglin T. R., 1996 Warthog and groundhog, novel families related to hedgehog. Curr. Biol. 6: 1047–1050.

Bürglin T. R., and P. E. Kuwabara, 2006 Homologs of the Hh signalling network in C. elegans. WormBook 1–14. 10.1895/wormbook.1.76.1

Bürglin T. R., 2008 Evolution of hedgehog and hedgehog-related genes, their origin from Hog proteins in ancestral eukaryotes and discovery of a novel Hint motif. BMC Genomics 9: 1–28. 10.1186/1471-2164-9-127

Burkhart D. L., and J. Sage, 2008 Cellular mechanisms of tumour suppression by the retinoblastoma gene. Nat. Rev. Cancer 8: 671–682. 10.1038/nrc2399

Cao J., J. S. Packer, V. Ramani, D. A. Cusanovich, C. Huynh, et al., 2017 Comprehensive single-cell transcriptional profiling of a multicellular organism. Science 357: 661–667. 10.1126/science.aam8940

Chen J., L. Y. Tang, M. E. Powell, J. M. Jordan, and L. R. Baugh, 2022 Genetic analysis of daf-18/PTEN missense mutants for starvation resistance and developmental regulation during Caenorhabditis elegans L1 arrest. G3 Genes, Genomes, Genet. 12. 10.1093/g3journal/jkac092

Chen J., R. Chitrakar, and R. Baugh, 2025 DAF-18/PTEN protects LIN-35/Rb from CLP-1/CAPN-mediated cleavage to promote starvation resistance. bioRxiv 2025.02.17.638677. 10.1101/2025.02.17.638677

Cochrane C. R., V. Vaghjiani, A. Szczepny, W. S. N. Jayasekara, A. Gonzalez-Rajal, et al., 2020 Trp53 and Rb1 regulate autophagy and ligand-dependent Hedgehog signaling. J. Clin. Invest. 140: 4006–4018. 10.1172/JCI132513

Dorman J. B., B. Albinder, T. Shroyer, and C. Kenyon, 1995 The age-1 and daf-2 genes function in a common pathway to control the lifespan of Caenorhabditis elegans. Genetics 141: 1399–1406. 10.1093/genetics/141.4.1399

Espelt M. V, A. Y. Estevez, X. Yin, and K. Strange, 2005 Oscillatory Ca2+ signaling in the isolated Caenorhabditis elegans intestine: role of the inositol-1,4,5-trisphosphate receptor and phospholipases C beta and gamma. J. Gen. Physiol. 126: 379–392. 10.1085/jgp.200509355

Falsztyn I. B., S. M. Taylor, and L. R. Baugh, 2025 Developmental and conditional regulation of DAF-2/INSR ubiquitination in Caenorhabditis elegans. G3 (Bethesda). 10.1093/g3journal/jkaf009

Fletcher M., E. J. Tillman, V. L. Butty, S. S. Levine, and D. H. Kim, 2019 Global transcriptional regulation of innate immunity by ATF-7 in C. elegans. PLoS Genet. 15: 1–14. 10.1371/journal.pgen.1007830

Fukuyama M., A. E. Rougvie, and J. H. Rothman, 2006 C. elegans DAF-18/PTEN mediates nutrient-dependent arrest of cell cycle and growth in the germline. Curr. Biol. 16: 773–779. 10.1016/j.cub.2006.02.073

Fukuyama M., K. Sakuma, R. Park, H. Kasuga, R. Nagaya, et al., 2012 C. Elegans AMPKs promote survival and arrest germline development during nutrient stress. Biol. Open 1:929–936. 10.1242/bio.2012836

Godfrey K. M., and D. J. Barker, 2001 Fetal programming and adult health. Public Health Nutr. 4: 611–624. 10.1079/phn2001145

Govindan J. A., E. Jayamani, X. Zhang, P. Breen, J. Larkins-Ford, et al., 2015 Lipid signalling couples translational surveillance to systemic detoxification in Caenorhabditis elegans. Nat. Cell Biol. 17: 1294–1303. 10.1038/ncb3229

Han S. K., D. Lee, H. Lee, D. Kim, H. G. Son, et al., 2016 OASIS 2: Online application for survival analysis 2 with features for the analysis of maximal lifespan and healthspan in aging research. Oncotarget 7: 56147–56152. 10.18632/oncotarget.11269

Hao L., K. Mukherjee, S. Liegeois, D. Baillie, M. Labouesse, et al., 2006 The hedgehog-related gene qua-1 is required for molting in Caenorhabditis elegans. Dev. Dyn. 235: 1469–1481. 10.1002/dvdy.20721

Hibshman J. D., A. E. Doan, B. T. Moore, R. E. Kaplan, A. Hung, et al., 2017 daf-16/FoxO promotes gluconeogenesis and trehalose synthesis during starvation to support survival. Elife 6: 1–29. 10.7554/eLife.30057

Holdorf A. D., D. P. Higgins, A. C. Hart, P. R. Boag, G. J. Pazour, et al., 2020 WormCat: An online tool for annotation and visualization of caenorhabditis elegans genome-scale data. Genetics 214: 279–294. 10.1534/genetics.119.302919

Hoppe T., G. Cassata, J. M. Barral, W. Springer, A. H. Hutagalung, et al., 2004 Regulation of the myosin-directed chaperone UNC-45 by a novel E3/E4-multiubiquitylation complex in C. elegans. Cell 118: 337–349. 10.1016/j.cell.2004.07.014

Hu Q., D. R. D’Amora, L. T. MacNeil, A. J. M. Walhout, and T. J. Kubiseski, 2017 The Oxidative Stress Response in Caenorhabditis elegans Requires the GATA Transcription Factor ELT-3 and SKN-1/Nrf2. Genetics 206: 1909–1922. 10.1534/genetics.116.198788

Jiang J., and C.-C. Hui, 2008 Hedgehog signaling in development and cancer. Dev. Cell 15: 801–812. 10.1016/j.devcel.2008.11.010

Jobson M. A., J. M. Jordan, M. A. Sandrof, J. D. Hibshman, A. L. Lennox, et al., 2015 Transgenerational effects of early life starvation on growth, reproduction, and stress resistance in Caenorhabditis elegans. Genetics 201: 201–212. 10.1534/genetics.115.178699

Jordan J. M., J. D. Hibshman, A. K. Webster, R. E. W. Kaplan, A. Leinroth, et al., 2019 Insulin/IGF Signaling and Vitellogenin Provisioning Mediate Intergenerational Adaptation to Nutrient Stress. Curr. Biol. 29: 2380–2388.e5. 10.1016/j.cub.2019.05.062

Jordan J. M., A. K. Webster, J. Chen, R. Chitrakar, and L. Ryan Baugh, 2023 Early-life starvation alters lipid metabolism in adults to cause developmental pathology in Caenorhabditis elegans. Genetics 223: 1–12. 10.1093/genetics/iyac172

Kaplan R. E. W., Y. Chen, B. T. Moore, J. M. Jordan, C. S. Maxwell, et al., 2015 dbl-1/TGF-β and daf-12/NHR Signaling Mediate Cell-Nonautonomous Effects of daf-16/FOXO on Starvation-Induced Developmental Arrest. PLoS Genet. 11: 1–23. 10.1371/journal.pgen.1005731

Kumsta C., and M. Hansen, 2012 C. elegans rrf-1 mutations maintain RNAi efficiency in the soma in addition to the germline. PLoS One 7: 1–12. 10.1371/journal.pone.0035428

Kuwabara P. E., M. H. Lee, T. Schedl, and G. S. X. E. Jefferis, 2000 A C. elegans patched gene, ptc-1, functions in germ-line cytokinesis. Genes Dev. 14: 1933–1944. 10.1101/gad.14.15.1933

Latorre I., M. A. Chesney, J. M. Garrigues, P. Stempor, A. Appert, et al., 2015 The DREAM complex promotes gene body H2A.Z for target repression. Genes Dev. 29: 495–500. 10.1101/gad.255810.114

Lee Y. R., M. Chen, and P. P. Pandolfi, 2018 The functions and regulation of the PTEN tumour suppressor: new modes and prospects. Nat. Rev. Mol. Cell Biol. 19: 547–562. 10.1038/s41580-018-0015-0

Lee Y., Y. Jung, D.-E. Jeong, W. Hwang, S. Ham, et al., 2021 Reduced insulin/IGF1 signaling prevents immune aging via ZIP-10/bZIP-mediated feedforward loop. J. Cell Biol. 220. 10.1083/jcb.202006174

Li J., C. Yen, D. Liaw, K. Podsypanina, S. Bose, et al., 1997 PTEN, a putative protein tyrosine phosphatase gene mutated in human brain, breast, and prostate cancer. Science 275: 1943–1947. 10.1126/science.275.5308.1943

Liu K., M. Grover, F. Trusch, C. Vagena-Pantoula, D. Ippolito, et al., 2024 Paired C-type lectin receptors mediate specific recognition of divergent oomycete pathogens in C. elegans. Cell Rep. 43: 114906. 10.1016/j.celrep.2024.114906

Lu X., and H. Robert Horvitz, 1998 lin-35 and lin-53, Two Genes that Antagonize a C. elegans Ras Pathway, Encode Proteins Similar to Rb and Its Binding Protein RbAp48 the activities of the ETS transcription factor LIN-1 and the winged-helix transcription factor LIN-31 (reviewed by Horvitz . Cell 95: 981–991.

Magliano M. P. Di, and M. Hebrok, 2003 Hedgehog signalling in cancer formation and maintenance. Nat. Rev. Cancer 3: 903–911. 10.1038/nrc1229

Marré J., E. C. Traver, and A. M. Jose, 2016 Extracellular RNA is transported from one generation to the next in Caenorhabditis elegans. Proc. Natl. Acad. Sci. U. S. A. 113: 12496–12501. 10.1073/pnas.1608959113

Mathies L. D., M. Schvarzstein, K. M. Morphy, R. Blelloch, A. M. Spence, et al., 2004 TRA-1/GLI controls development of somatic gonadal precursors in C. elegans. Development 131: 4333–4343. 10.1242/dev.01288

Mitani S., 2017 Comprehensive functional genomics using Caenorhabditis elegans as a model organism. Proc. Jpn. Acad. Ser. B. Phys. Biol. Sci. 93: 561–577. 10.2183/pjab.93.036

Morris J. Z., H. A. Tissenbaum, and G. Ruvkun, 1996 A phosphatidylinositol-3-OH kinase family member regulating longevity and diapause in Caenorhabditis elegans. Nature 382: 536–539. 10.1038/382536a0

Muñoz M. J., and D. L. Riddle, 2003 Positive selection of Caenorhabditis elegans mutants with increased stress resistance and longevity. Genetics 163: 171–180. 10.1093/genetics/163.1.171

Murphy C. T., and P. J. Hu, 2013 Insulin/insulin-like growth factor signaling in C. elegans. WormBook 1–43. 10.1895/wormbook.1.164.1

Myers M. P., J. P. Stolarov, C. Eng, J. Li, S. I. Wang, et al., 1997 P-TEN, the tumor suppressor from human chromosome 10q23, is a dual-specificity phosphatase. Proc. Natl. Acad. Sci. U. S. A. 94: 9052–9057. 10.1073/pnas.94.17.9052

Myers M. P., I. Pass, I. H. Batty, J. Van der Kaay, J. P. Stolarov, et al., 1998 The lipid phosphatase activity of PTEN is critical for its tumor supressor function. Proc. Natl. Acad. Sci. U. S. A. 95: 13513–13518. 10.1073/pnas.95.23.13513

Nüsslein-Volhard C., and E. Wieschaus, 1980 Mutations affecting segment number and polarity in Drosophila. Nature 287: 795–801. 10.1038/287795a0

Ogg S., and G. Ruvkun, 1998 The C. elegans PTEN Homolog, DAF-18, Acts in the Insulin Receptor-like Metabolic Signaling Pathway is a member of the insu-lin/insulin-like growth factor family of receptors (Kimura et al., 1997). Activation of the DAF-2 signaling cascade is necessary for. Mol. Cell 2: 887–893.

Olmedo M., A. Mata-Cabana, M. Jesús Rodríguez-Palero, S. García-Sánchez, A. Fernández-Yañez, et al., 2020 Prolonged quiescence delays somatic stem cell-like divisions in Caenorhabditis elegans and is controlled by insulin signaling. Aging Cell 19: 1–13. 10.1111/acel.13085

Paek J., J. Y. Lo, S. D. Narasimhan, T. N. Nguyen, K. Glover-Cutter, et al., 2012 Mitochondrial SKN-1/Nrf mediates a conserved starvation response. Cell Metab. 16: 526–537. 10.1016/j.cmet.2012.09.007

Qadota H., M. Inoue, T. Hikita, M. Köppen, J. D. Hardin, et al., 2007 Establishment of a tissue-specific RNAi system in C. elegans. Gene 400: 166–173. 10.1016/j.gene.2007.06.020

Ravelli G. P., Z. A. Stein, and M. W. Susser, 1976 Obesity in young men after famine exposure in utero and early infancy. N. Engl. J. Med. 295: 349–353. 10.1056/NEJM197608122950701

Rohlfing A.-K., Y. Miteva, L. Moronetti, L. He, and T. Lamitina, 2011 The Caenorhabditis elegans mucin-like protein OSM-8 negatively regulates osmosensitive physiology via the transmembrane protein PTR-23. PLoS Genet. 7: e1001267. 10.1371/journal.pgen.1001267

Shaul N. C., J. M. Jordan, I. B. Falsztyn, and L. Ryan Baugh, 2023 Insulin/IGF-dependent Wnt signaling promotes formation of germline tumors and other developmental abnormalities following early-life starvation in Caenorhabditis elegans. Genetics 223: 1–11. 10.1093/genetics/iyac173

Sherry T., H. Nicholas, and R. Pocock, 2019 New deletion alleles for Caenorhabditis elegans Hedgehog pathway-related genes wrt-6 and wrt-10. microPublication Biol. 2019. 10.17912/micropub.biology.000169

Shivers R. P., M. J. Youngman, and D. H. Kim, 2008 Transcriptional responses to pathogens in Caenorhabditis elegans. Curr. Opin. Microbiol. 11: 251–256. 10.1016/j.mib.2008.05.014

Shivers R. P., T. Kooistra, S. W. Chu, D. J. Pagano, and D. H. Kim, 2009 Tissue-Specific Activities of an Immune Signaling Module Regulate Physiological Responses to Pathogenic and Nutritional Bacteria in C. elegans. Cell Host Microbe 6: 321–330. 10.1016/j.chom.2009.09.001

Sigafoos A. N., B. D. Paradise, and M. E. Fernandez-Zapico, 2021 Hedgehog/gli signaling pathway: Transduction, regulation, and implications for disease. Cancers (Basel). 13. 10.3390/cancers13143410

Sijen T., J. Fleenor, F. Simmer, K. L. Thijssen, S. Parrish, et al., 2001 On the role of RNA amplification in dsRNA-triggered gene silencing. Cell 107: 465–476. 10.1016/s0092-8674(01)00576-1

Steck P. A., M. A. Pershouse, S. A. Jasser, W. K. Yung, H. Lin, et al., 1997 Identification of a candidate tumour suppressor gene, MMAC1, at chromosome 10q23.3 that is mutated in multiple advanced cancers. Nat. Genet. 15: 356–362. 10.1038/ng0497-356

Sternberg P. W., K. Van Auken, Q. Wang, A. Wright, K. Yook, et al., 2024 WormBase 2024: status and transitioning to Alliance infrastructure. Genetics 227: 1–14. 10.1093/genetics/iyae050

Stiernagle T., 2006 Maintenance of C. elegans. WormBook 1–11. 10.1895/wormbook.1.101.1

Tabara H., M. Sarkissian, W. G. Kelly, J. Fleenor, A. Grishok, et al., 1999 The rde-1 gene, RNA interference, and transposon silencing in C. elegans. Cell 99: 123–132. 10.1016/S0092-8674(00)81644-X

Tan M. W., S. Mahajan-Miklos, and F. M. Ausubel, 1999 Killing of Caenorhabditis elegans by Pseudomonas aeruginosa used to model mammalian bacterial pathogenesis. Proc. Natl. Acad. Sci. U. S. A. 96: 715–720. 10.1073/pnas.96.2.715

Tawo R., W. Pokrzywa, É. Kevei, M. E. Akyuz, V. Balaji, et al., 2017 The Ubiquitin Ligase CHIP Integrates Proteostasis and Aging by Regulation of Insulin Receptor Turnover. Cell 169: 470–482.e13. 10.1016/j.cell.2017.04.003

Templeman N. M., V. Cota, W. Keyes, R. Kaletsky, and C. T. Murphy, 2020 CREB Non-autonomously Controls Reproductive Aging through Hedgehog/Patched Signaling. Dev. Cell 54: 92–105.e5. 10.1016/j.devcel.2020.05.023

Tepper R. G., J. Ashraf, R. Kaletsky, G. Kleemann, C. T. Murphy, et al., 2013 PQM-1 complements DAF-16 as a key transcriptional regulator of DAF-2-mediated development and longevity. Cell 154: 676–690. 10.1016/j.cell.2013.07.006

Tijsterman M., J. Pothof, and R. H. A. Plasterk, 2002 Frequent germline mutations and somatic repeat instability in DNA mismatch-repair-deficient Caenorhabditis elegans. Genetics 161: 651–660. 10.1093/genetics/161.2.651

VanDerMolen K. R., M. A. Newman, P. C. Breen, Y. Gao, L. A. Huff, et al., 2025 Non-cell-autonomous regulation of mTORC2 by Hedgehog signaling maintains lipid homeostasis. Cell Rep. 44: 115191. 10.1016/j.celrep.2024.115191

Vélez-Cruz R., and D. G. Johnson, 2017 The retinoblastoma (RB) tumor suppressor: Pushing back against genome instability on multiple fronts. Int. J. Mol. Sci. 18. 10.3390/ijms18081776

Walker C. L., and S. M. Ho, 2012 Developmental reprogramming of cancer susceptibility. Nat. Rev. Cancer 12: 479–486. 10.1038/nrc3220

Wang Q., R. Fu, G. Li, S. Xiong, Y. Zhu, et al., 2023 Hedgehog receptors exert immune-surveillance roles in the epidermis across species. Cell Rep. 42: 112929. 10.1016/j.celrep.2023.112929

Wolf T., W. Qi, V. Schindler, E. D. Runkel, and R. Baumeister, 2014 Doxycyclin ameliorates a starvation-induced germline tumor in C. elegans daf-18/PTEN mutant background. Exp. Gerontol. 56: 114–122. 10.1016/j.exger.2014.04.002

Yen K., S. D. Narasimhan, and H. A. Tissenbaum, 2011 DAF-16/forkhead Box O transcription factor: Many paths to a single fork(head) in the road. Antioxidants Redox Signal. 14: 623– 634. 10.1089/ars.2010.3490

Zárate-Potes A., I. Ali, M. Ribeiro Camacho, H. Brownless, and A. Benedetto, 2022 Meta-Analysis of Caenorhabditis elegans Transcriptomics Implicates Hedgehog-Like Signaling in Host-Microbe Interactions. Front. Microbiol. 13: 1–17. 10.3389/fmicb.2022.853629

Zhang Y., and P. A. Beachy, 2023 Cellular and molecular mechanisms of Hedgehog signalling. Nat. Rev. Mol. Cell Biol. 24: 668–687. 10.1038/s41580-023-00591-1

Zugasti O., J. Rajan, and P. E. Kuwabara, 2005 The function and expansion of the Patchedand Hedgehog-related homologs in C. elegans. Genome Res. 15: 1402–1410. 10.1101/gr.3935405

